# Scoring alignments by embedding vector similarity

**DOI:** 10.1101/2023.08.30.555602

**Authors:** Sepehr Ashrafzadeh, G. Brian Golding, Silvana Ilie, Lucian Ilie

## Abstract

Sequence similarity is of paramount importance in biology, as similar sequences tend to have similar function and share common ancestry. Scoring matrices, such as PAM or BLO-SUM, play a crucial role in all bioinformatics algorithms for identifying similarities, but have the drawback that they are fixed, independent of context. We propose a new scoring method for amino acid similarity that remedies this weakness, being contextually dependent. It relies on recent advances in deep learning architectures that employ self-supervised learning in order to leverage the power of enormous amounts of unlabelled data to generate contextual embeddings, which are vector representations for words. These ideas have been applied to protein sequences, producing embedding vectors for protein residues. We propose the *E-score* between two residues as the cosine similarity between their embedding vector representations. Thorough testing on a wide variety of reference multiple sequence alignments indicate that the alignments produced using the new *E*-score method, especially ProtT5-score, are significantly better than those obtained using BLOSUM matrices. The new method proposes to change the way alignments are computed, with far reaching implications in all areas of textual data that use sequence similarity. The program to compute alignments based on various *E*-scores is available as a web server at e-score.csd.uwo.ca. The source code is freely available for download from github.com/lucian-ilie/E-score.

## 1 Introduction

Sequence similarity is fundamental in sequence analysis. Significant similarities do not appear by chance but are evolutionarily motivated, with corresponding implications on their associated functions. Given a new sequence, its functionality is much faster investigated by performing a database search to identify known similar sequences and inferring the function of the new sequence from the existing information, resulting often in accurate identification. This procedure is sometimes the only way to infer function. Sequence similarity search is the most widely used procedure in bioinformatics, the BLAST program [2, 3] being one of the most cited contributions in the history of science.^1^

The most important component of similarity search algorithms, such as BLAST, is the function that describes the similarity scores of various components, such as nucleotides for DNA and amino acids for proteins. We consider here the much more complicated case of proteins, where scoring of alignments is done using PAM [6] or BLOSUM [11] matrices, which give the substitution scores for pairs of amino acids. These matrices are fixed, which means that the substitution score for a pair of amino acids does not depend on their context, that is, the surrounding sequence of the protein containing them. We propose a new scoring function for residue pairs that is dependent on the context in which the residues appear. The new score makes use of protein embeddings.

Many of the best current techniques in Natural Language Processing (NLP) involve transfer learning based on pretrained models using self-supervised learning on unlabelled data. Such data is freely available in enormous amounts, such as the Common Crawl corpus (commoncrawl.org). Words that share similar meanings or are related often appear in similar contexts. This enables scalable models to map words to numerical vectors (embeddings) in a high-dimensional space. Words with similar co-occurrence patterns are represented by vectors that are closer in this space. Such text embeddings are able to capture the semantic meaning of words. One of the best known earlier models is word2vec [21]. Denoting by *w*_*v*_ the vector associated with a word *w*, an example ([21]) of meaning capture is shown by the fact that `king`_*v*_ − `man`_*v*_ + `woman`_*v*_ produces a vector that is closest to `queen`_*v*_.

Embeddings such as word2vec [21] or GloVe [24] are contextual independent as the embedding vector for a given word is the same regardless of context. The next generation of embeddings, called contextual embeddings, overcome this limitation, e.g., ELMo [25], BERT [7], RoBERTa [19], XLNet [34], T5 [26]. The most successful contextual embeddings rely on the transformer architecture [32], achieving state-of-the-art results in many NLP benchmarks [7, 19, 34, 18].

Protein sequences present a similar situation, with entire proteins seen as sentences and residues as words, enabling the use of the embedding techniques developed for NLP to take advantage of the vast amount of protein sequences from UniProt [1]. Many protein embeddings have been proposed, including ProtVec [4], SeqVec [10], PRoBERTa [22], MSA-transformer [27], ESM-1b and ESM2 [28], and the models in the ProtTrans project: ProtTXL, ProtBert, ProtXLNet, ProtAlbert, ProtElectra, and ProtT5 [8]. These protein embeddings have many important applications to various areas, such as: structure prediction [29, 33, 15], function prediction [16, 9, 17], interaction site prediction [13, 35, 12], etc. In many applications they help achieving state-of-the-art performance.

Our new scoring function uses any embedding method that associates a vector to each residue. For any such method, *E*, we define the *E-score* between two residues as the cosine similarity between their associated vectors. We investigate this new scoring function for contextual embeddings from the point of view of correct alignment generation. We have selected a wide variety of reference multiple sequence alignments (MSAs) from NCBI’s CDD (Conserved Domain Database) [20], that are manually curated according to the function. For each, we randomly selected many pairs of protein sequences in order to check how close the pairwise alignment is to the original reference MSA. Several alignment distances [5], including two that we introduce here, were used to compare the best new contextual ProtT5-score, with the best old one, the BLOSUM45 matrix. ProtT5-score performs overwhelmingly better than BLOSUM45, producing an average alignment distance that is better in all but one of the 38 MSAs tested.

The new scoring method proposes completely changing the way alignments are computed, with far reaching implications in all areas that use sequence similarity.

## 2 Materials and Methods

### 2.1 Cosine similarity

For two vectors, *A* = (*A*_*i*_)_*i*=1..*n*_ and *B* = (*B*_*i*_)_*i*=1..*n*_, the *cosine similarity* between *A* and *B* is defined as the cosine of the angle, *θ*, between the two vectors:

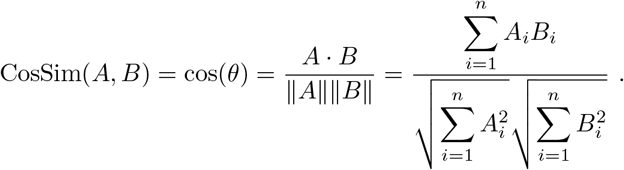

The cosine similarity takes values between −1 and −1, with 1 for vectors of opposite direction, 1 for the same direction, and 0 for orthogonal vectors; Fig. 1 illustrates these three cases.

**Figure 1.**
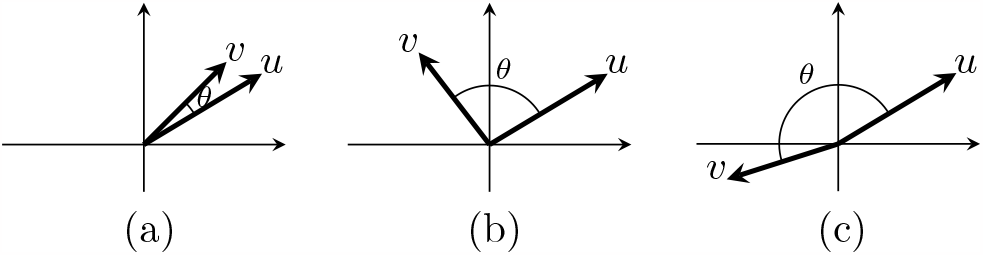
Cosine similarity of vectors *u* and *v* is the cosine of the angle, *θ*, between them: (a) similar vectors: *θ* ≈ 0, cos(*θ*) ≈ 1; (b) orthogonal (independent) vectors: 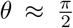, cos(*θ*) ≈ 0; (c) opposite vectors: *θ* ≈ *π*, cos(*θ*) ≈ −1.

### 2.2 E-score

For a protein sequence *P*, of length *m*, denote its *i*^th^ residue by *P* [*i*]. Consider an embedding method, *E*, that produces embedding vectors of size *n*. Assume that *E*, when applied to protein *P*, generates an *m × n* matrix, *E*(*P*), that has as its rows all *m* embedding vectors of the residues of *P* ; denote the embedding of the *i*^th^ residue by *E*(*P*)_*i*_ = *E*(*P*)(*i*, :).

For two proteins *P* and *Q*, and two positions, *i* within *P* and *j* within *Q*, the *E-score* between the residues *P* [*i*] and *Q*[*j*] is defined as the cosine similarity between the corresponding embedding vectors:

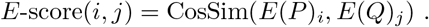

Note that, while not reflected by our notation, *E*-score(*i, j*) depends also on the protein sequences *P* and *Q*. Thus, when the embedding method is ProtT5, ProtBert, ESM2, etc., we obtain ProtT5-score, ProtBert-score, ESM2-score, and so on. Clearly, for any contextual embedding method *E*, the *E*-score changes with context as well.

### 2.3 Visualizing *E*-scores

To have an idea what these new *E*-scores look like, we constructed some scoring matrices using those, and compared them with the BLOSUM matrices. While a BLOSUM matrix is fixed, the *E*-scores change with context, so we computed average scores for each pair of amino acids. We picked the longest multiple sequence alignment (MSA), *NBD_sugar-kinase_HSP70_actin*, from the selected six representative MSAs (from the first column of Table 3). Then, we distinguished between “aligned” residues, which appear in the same column of the MSA, and “unaligned” residues, which occur in different columns. Since the *E*-scores use the context, it is expected that aligned residues have significantly higher scores compared to unaligned ones. For each pair of amino acids, we randomly selected occurrences of their residues in the MSA and computed the average score; 100,000 pairs were selected in each of the categories, aligned and unaligned. We do this separately for the aligned and the unaligned case. This way, for a given embedding type and an MSA, we obtain two 20 *×* 20 matrices. We show the aligned matrices for three embeddings: ProtT5-score, ESM2-score, and ProtAlbert-score, alongside the BLOSUM45 matrix, in Fig. 2. It appears from Fig. 2 that the matrix for the ProtT5-score is the most similar to the BLOSUM45 matrix. The heatmaps of all five BLOSUM matrices and all twelve *E*-score matrices (two matrices for each embedding method, for the aligned and the unaligned case) are shown in the Supplementary Table 8. It is interesting to notice the wide variety of ranges among the *E*-score matrices. The unaligned matrices have lower, and narrower, ranges. It appears that a wider range of values is an indication of better performance for alignments.

**Figure 2.**
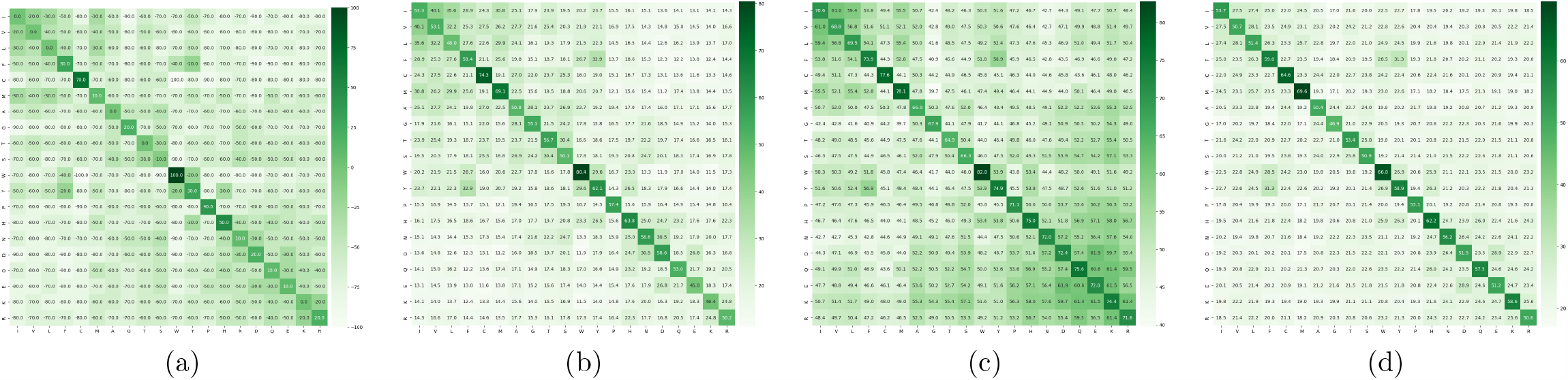
Heatmaps of (a) BLOSUM45 matrix (scaled to 1 −1) and three aligned matrices of average *E*-scores for the *NBD_sugar-kinase_HSP70_actin* MSA: (b) ProtT5-score, (c) ESM2-score, and (d) ProtAlbert-score.

### 2.4 Correlations

It is of interest to see how well the new *E*-scores are correlated with the BLOSUM matrices. We consider here five BLOSUM matrices: BLOSUM45, BLOSUM50, BLOSUM62, BLOSUM80, and BLOSUM90, and six embedding methods: ProtT5, ProtBert, ProtAlbert, ProtXLNet, ESM-1b and ESM2. For each embedding method, we use for computing the correlations the same matrices computed above for the *NBD_sugar-kinase_HSP70_actin* MSA. Figure 3 gives the Pearson correlations between the five BLOSUM matrices and the twelve *E*-score matrices. It can be seen that the ProtT5 matrices have the highest correlation with all BLOSUM matrices, followed, in order, by ESM2 and ProtBert, and then ProtAlbert, ESM-1b and, by far the last, ProtXLNet. For each method, the aligned matrix has higher correlation with BLOSUM matrices than the unaligned one. Interestingly, the unaligned ProtT5 matrix comes in second place, after the aligned ProtT5 and before all the other aligned matrices. With the exception of ProtXLNet, the two matrices, aligned and unaligned, for the same embedding method, are highly correlated.

**Figure 3.**
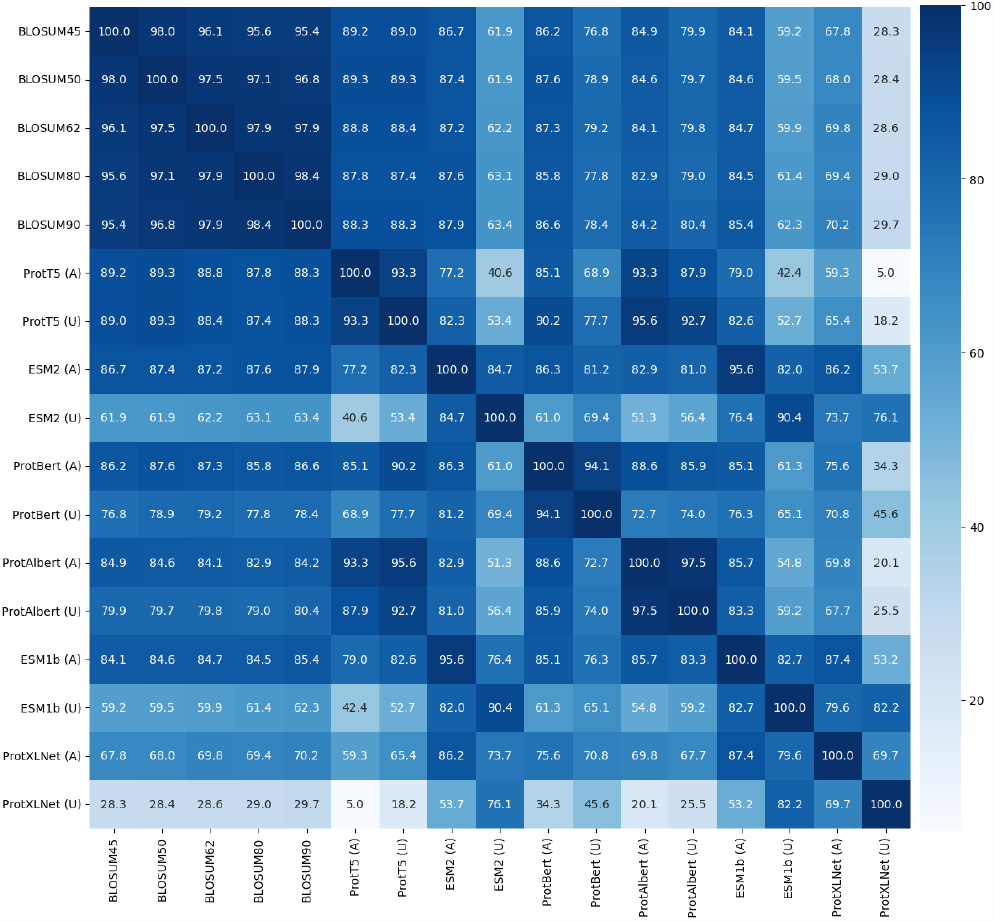
Pearson correlations between five BLOSUM matrices and twelve *E*-score matrices. For each embedding method we considered, the aligned and unaligned matrices for the *NBD_sugar-kinase_HSP70_actin* MSA have been included.

### 2.5 Alignment distances

Our goal is to show that the use of the new *E*-scores provides better alignments compared to the BLOSUM scores. We do this by comparing their ability to produce true alignments, as given by the reference alignments from CDD; see Reference alignments subsection below. We selected a number of reference MSAs, discussed in the Results section, and investigated how close pairwise alignments produced using *E*-score and BLOSUM matrices, resp., are from the alignments between the same sequences, as induced by the reference MSAs. To compare how close two alignments are, we need to measure distances between alignments. We use five distance metrics, three existing ones, d_ssp_, d_seq_, and d_pos_ from [5], and two that we introduce here, d_cc_ and d_d_.

We give first a formal definition of an alignment. Consider an alphabet Σ. For a string *P* over Σ of length |*P* | = *n*, let *P* [*i*], 1 ≤ *i* ≤ *n*, denote the *i*th character of *P* ; thus *P* = *P* [1]*P* [2] *· · · P* [*n*]. For two strings *P, Q* over Σ, an *alignment* between *P* and *Q* is a pair of strings *A* = (*P*_*A*_, *Q*_*A*_), of the same length |*P*_*A*_| = |*Q*_*A*_|, over Σ ∪ *{*–*}* (the ‘ –’ character stands for a *gap*), with *P*_*A*_ (*Q*_*A*_) being obtained by inserting gaps at any positions in *P* (*Q*, resp.), with the restriction that, for any position *i*, at least one of the characters *P*_*A*_[*i*] and *Q*_*A*_[*i*] is not a gap.

**Example 1**. Here are two examples of alignments between peptides *P* = `RKD` and *Q* = `DNN`:

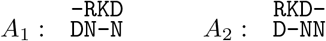

We have, 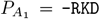 and 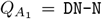, both of length 4. Note that there is no column with two gaps.

Four distances have been introduced by [5], of which we use three that are suitable for our purpose: d_ssp_, symmetrized sum-of-pairs score, that ignores gaps; d_seq_, that treats all gaps in the same sequence equally; and d_pos_, that incorporates positional information about the gaps. The original alignment is modified to reflect each gap treatment; in Example 1, alignment *A*_1_ is converted into the following three alignments, to be used for each of the three distances d_ssp_, d_seq_, d_pos_, respectively:

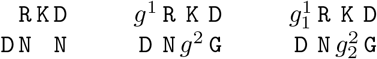

Similarly, *A*_2_ is converted into the following three versions:

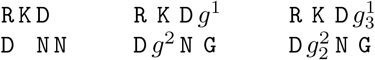

The three distances, d_ssp_, d_seq_, d_pos_, are then defined using the three versions above, in order, using the homology set for each character, 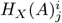, for alignment *A, X*∈ {ssp, seq, pos}, sequence *i*, position *j*, that contain the characters from the other sequences in the same column as *j*. For example, 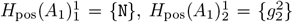. d_ssp_ is defined as the Jaccard distance [14] on the homology sets *H*_ssp_:

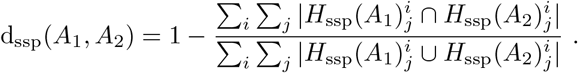

The other two distances use symmetric difference on the same sets; for *X* ∈ *{*seq, pos*}*:

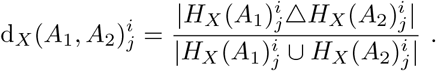

We refer the reader to [5] for details and examples.

### 2.6 Relative displacement distance

The fourth distance, d_d_, is new and measures the relative displacement between the positions of the letters in the two strings. Put |*P* | = *n* and |*Q*| = *m* and let 1 ≤ *p*_1_ *< p*_2_ *< · · · < p*_*n*_ ≤ |*P*_*A*_| be such that *P*_*A*_[*p*_*i*_] = *P* [*i*], for any 1 ≤ *i* ≤ *n*. Similarly, let *q*_1_ *< q*_2_ *< · · · < q*_*m*_ be such that *Q*_*A*_[*q*_*j*_] = *Q*[*j*], for any 1 ≤*j* ≤*m*. For any *i*, 1 ≤*i*≤ *n*, and *j*, 1 ≤*j* ≤*m*, denote the distance between the positions of *P* [*i*] and *Q*[*j*] in *A* by *d*_*A*_(*i, j*) = *p*_*i*_ − *q*_*j*_.

Given two alignments, *A*_1_ and *A*_2_, between the same strings, we define the *relative displacement distance* between them as:

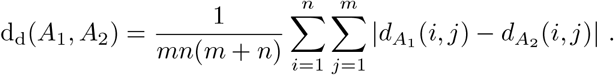

The division by *mn*(*m* + *n*) is done to scale the value between 0 and 1. For any *i, j* with 1 ≤ *i* ≤ *n*, 1 ≤ *j* ≤ *m*, we have that *i* ≤ *p*_*i*_ ≤ *m* + *i* and *j* ≤ *q*_*j*_ ≤ *n* + *j*, implying that *i*−*n*−*j* ≤ *p*_*i*_−*q*_*j*_ ≤ *m*+*i*−*j*. Therefore, 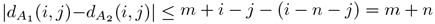.

Example 1: For alignment *A*_1_, we have *n* = *m* = 3, and (*p*_1_, *p*_2_, *p*_3_) = (2, 3, 4), (*q*_1_, *q*_2_, *q*_3_) = (1, 2, 4). For *A*_2_, we have (*p*_1_, *p*_2_, *p*_3_) = (1, 2, 3) and (*q*_1_, *q*_2_, *q*_3_) = (1, 3, 4). Then, 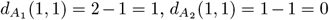. The distance between the alignments is 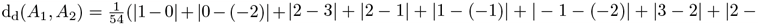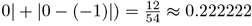.

### 2.7 Closest context distance

The first three distances above, d_ssp_, d_seq_, and d_pos_, consider only the column context of each residue, disregarding the position at which this occurs. On the other hand, the d_d_ distance considers only the relative positions of the residues, disregarding their contexts. We propose next a distance that captures both position and context, by considering the distance to the closest position with the same context.

We introduce more notation. For a string *P* over Σ, of length |*P*| = *n*, denote the range *P* [*i*..*j*] = *P* [*i* + 1]*P* [*i* + 2] *· · · P* [*j*], if *i* ≤*j*, and *P* [*i*..*j*] = *P* [*j*]*P* [*j* + 1] *· · · P* [*i* −1], if *i > j*. Note that *P* [*i, i*] = *ε*, the empty string.

For a string *w* over Σ ∪ *{* – *}*, denote by *ℒ* (*w*) the number of letters from Σ in *w*; e.g., *ℒ* (`DN-N`) = 3.

We modify slightly some notation from above. If |*P* | = *n* and |*Q*| = *m*, and let *p*_*ℓ,i*_ be the position in 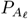 at which *P* [*i*] occurs; that is, 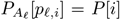, for any 1 ≤ *i* ≤ *n* and *f* = 1, 2. Similarly, let *q*_*ℓ,j*_ be the position in 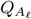 at which *Q*[*j*] occurs; that is, 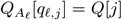, for any 1 ≤ *j* ≤ *m* and *ℓ* = 1, 2.

For a given *i*, 1 ≤ *i* ≤ *n*, we define *d*_*P,i*,1_, the contribution of *P* [*i*] from alignment *A*_1_ to the distance, as follows, depending on whether the letter opposite to 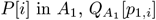, is (a) a letter or (b) a gap.

(a) If 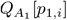 is a letter, then *d*_*P,i*,1_ is the number of letters in 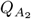 from position *p*_2,*i*_ to the closest occurrence of a residue that is the same as 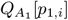 (“closest” means fewest letters; gaps are not counted):

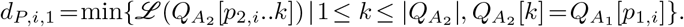 In Example 1, for *i* = 1, we have *p*_1,1_ = 2, *p*_2,1_ = 1, 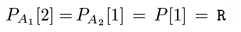 and 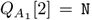. The closest `N` to the position 1 in 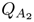 is at position 3, and there is one letter, `D`, in the range from 1 to 3. Therefore, 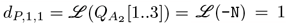. For *i* = 3, we have 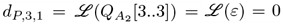, because *P* [3] = `D` has the same opposite residue, `N`, in both alignments.
(b) If 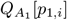 is a gap, then *d*_*P,i*,1_ is the difference between the number of letters in the two strings, 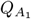 and 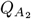, before the position of *P* [*i*] (note that this does not depend on whether 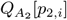 is a letter or a gap):

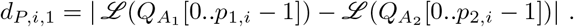 In Example 1, for *i* = 2, we have *p*_1,2_ = 3, *p*_2,2_ = 2, 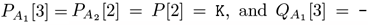 = –. Therefore, 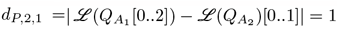.

The numbers *d*_*P,i*,2_, *d*_*Q,j*,1_, *d*_*Q,j*,2_, for 1 ≤ *i* ≤ *n*, 1 ≤ *j* ≤ *m*, are defined similarly. In our Example 1, we have 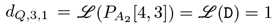.

Then, the *closest context distance* between *A*_1_ and *A*_2_ is:

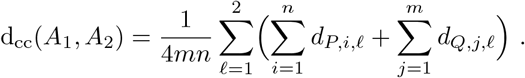

The division by 4*mn* is done to scale the value between 0 and 1, because we always have *d*_*P,i*, *ℓ*_ ≤ *m* and *d*_*Q,j*, *ℓ*_ ≤ *n*, thus making the sum after the fraction at most 4*mn*. In our Example 1, we have 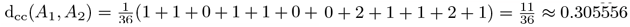.

## 3 Results

### 3.1 Reference alignments

There are multiple benchmark databases available to validate the accuracy of alignments. Most of these databases include comparatively similar sequences [31], albeit with the addition of features that are often difficult to align such as repeats, and many of the benchmarks were created using simulation techniques [30]. Most alignment algorithms can properly align similar sequences and the purpose of the method reported here is to align very distantly related protein sequences. For distantly related proteins we do not agree that simulation methods have completely captured the vagaries of protein sequence evolution. We therefore wanted to use very distantly related protein sequence alignments that had been done by hand and based on the function of the proteins. To this end, we choose to make use of the Conserved Domain Database (CDD), [20], from NCBI. This database has hand curated alignments and furthermore, it lists these alignments in a hierarchical fashion. Thus alignments can be chosen at different degrees of divergence from the same types of proteins.

Several alignments were chosen from CDD to explore different degrees of divergence and to explore different lengths of alignments. Pairs of protein groups were chosen to be very divergent (fold or superfamily level) and then a superfamily or family from within that protein group. Here are several examples: *Globin-like*, cd01067 (consensus alignment length 161 sites with 119 aa) and *Hb-alpha-like*, cd08927 (consensus alignment length 142 sites with 140 aa); *7tm_GPCRs*, cd14964, (consensus alignment length 420 sites with 267 aa) and *7tmA_photoreceptors insect*, cd15079 (consensus alignment length 301 sites with 292 aa); *NBD_sugarkinase_HSP70_actin*, cd00012 (consensus alignment length 1154 sites with 185 aa) and *FGGY_YpCarbK_like*, cd07782 (consensus alignment length 660 sites with 509 aa). The difference between the alignment length and the consensus number of aa gives an indication of the divergence within the grouping.

We have chosen 49 multiple sequence alignments from CDD. They are presented in Table 1, each with the conserved domain name, source, number of protein sequences, and alignment length. The list is sorted by alignment length.

**Table 1:** The 49 MSAs from the Conserved Domain Database: the columns give, in order, the conserved domain, source ID, number of protein sequences, and alignment length. The MSAs are sorted increasingly by length.

| MSAs |  |  |  |
| --- | --- | --- | --- |
| Conserved domain | Source | Proteins | Length |
| *Bbox2_MID2_C-I* | cd19823 | 8 | 40 |
| *Bbox2_TRIM42_C-III* | cd19782 | 9 | 40 |
| *Bbox2_MID* | cd19758 | 10 | 40 |
| *Bbox2_MID1_C-I* | cd19822 | 9 | 47 |
| *Bbox_SF* | cd00021 | 6 | 48 |
| *DEFL-defensin-like* | cd21806 | 78 | 51 |
| *Bbox2_TRIM37_C-VIII* | cd19779 | 25 | 52 |
| *Bbox2_TRIM9-like* | cd19764 | 17 | 53 |
| *CBD-like* | cd12204 | 41 | 61 |
| *Bbox2* | cd19756 | 127 | 65 |
| *ChtBD1* | cd00035 | 32 | 67 |
| *Bbox1_CYLD* | cd19816 | 27 | 68 |
| *KAZAL_FS* | cd00104 | 273 | 74 |
| *bHLH_SF* | cd00083 | 79 | 75 |
| *CD-CSD* | cd00024 | 522 | 98 |
| *C1* | cd00029 | 281 | 99 |
| *TrHb* | cd14756 | 8 | 130 |
| *Hb* | cd14765 | 15 | 138 |
| *SH2-STAT5* | cd10376 | 5 | 140 |
| *Hb-beta-like* | cd08925 | 27 | 140 |
| *Hb-alpha-like* | cd08927 | 39 | 142 |
| *SH2-STAT5a* | cd10421 | 10 | 145 |
| *Mb* | cd08926 | 9 | 149 |
| *GS-GGDEF-2* | cd14759 | 26 | 152 |
| *Globin-like* | cd01067 | 17 | 161 |
| *FHb-globin* | cd08922 | 47 | 162 |
| *PFM_HFR-2-like* | cd20216 | 90 | 174 |
| *SH2-STAT_family* | cd09919 | 67 | 206 |
| *PBP-like* | cd08919 | 31 | 213 |
| *SH2* | cd00173 | 352 | 214 |
| *Globin_sensor* | cd01068 | 193 | 223 |
| *PFM_monalysin-like* | cd17904 | 31 | 229 |
| *Mb-like* | cd01040 | 384 | 239 |
| *FYVE-like_SF* | cd00065 | 392 | 266 |
| *PFM_aerolysin_family* | cd10140 | 65 | 270 |
| *7tmA_photoreceptors_insect* | cd15079 | 35 | 301 |
| *7tmA_Melanopsin-like* | cd15083 | 12 | 314 |
| *ClyA_AhlB-like* | cd22652 | 39 | 354 |
| *7tmA_Opsins_type2_animals* | cd14969 | 71 | 400 |
| *7tm_GPCRs* | cd14964 | 18 | 420 |
| *7tmA_Anaphylatoxin_R-like* | cd14974 | 18 | 429 |
| *7tmA_Opioid_R-like* | cd14970 | 16 | 458 |
| *ClyA-like* | cd21116 | 118 | 519 |
| *FGGY_RBK-like* | cd07768 | 6 | 537 |
| *FGGY* | cd00366 | 31 | 588 |
| *FGGY_YpCarbK-like* | cd07782 | 35 | 690 |
| *7tm_classA_rhodopsin-like* | cd00637 | 405 | 808 |
| *NBD_sugar-kinase_HSP70_actin* | cd00012 | 125 | 1154 |
| *7tmA_amine_R-like* | cd14967 | 78 | 1227 |

### 3.2 Case studies

Before presenting a comparison between *E*-scores and the BLOSUM matrices, we show in this section two examples of the advantage presented by the alignments produced by the ProtT5-score compared with the BLOSUM45 matrix. The first example is for two sequences from the *KAZAL FS* (cd00104) MSA. The sequences and the two alignments are shown below. The BLOSUM45 alignment is correct in the first half, whereas the second half is significantly different, with several gaps added, whereas the true alignment has none.

– Sequence `S1`:

~~~
>gi|1827578|pdb|1TBR|R
CPHALHRVCGSDGETYSNPCTLNCAKFNGKPELVKVHDGPC
~~~

– Sequence `S2`:

~~~
>gi|124848|sp|P08480|IPSG_FELCA
CTMEYFPLCGSDGQEYSNKCLFCNEVVKRRGTLFLAKYGQC
~~~

–ProtT5-score end-gap-free alignment, which is identical with the reference MSA induced alignment:

~~~
S1 : 1 CPHALHRVCGSDGETYSNPCTLNCAKFNGKPELVKVHDGPC 41
       C       CGSDG  YSN C             L    G C
S2 : 1 CTMEYFPLCGSDGQEYSNKCLFCNEVVKRRGTLFLAKYGQC 41
~~~

–BLOSUM45 matrix end-gap-free alignment:

~~~
S1 : 1 CPHALHRVCGSDGETYSNPCTLNC---AKFNGKPELVKVHDGPC 41
       C      +CGSDG+ YSN C L C    K  G   L K   G C
S2 : 1 CTMEYFPLCGSDGQEYSNKC-LFCNEVVKRRGTLFLAKY--GQC 41
~~~

The second example involves two sequences from the *Globin sensor* (cd01068) MSA. The sequences and the two alignments are shown below. In this example, the BLOSUM45 alignment diverges drastically from the true alignment; the algorithm is deceived by a number of identities between the two sequences.

–Sequence `S1`:

~~~
>gi|374391926|gb|EHQ63256.1|
EQINYIGITDFDVDLLHSKETQFRAVVDMLVDELYEQITAQPELHRIILQHSTVERLKETQRWYFLSMAS
GVINEPFIEKRLHIGKVHSRIGLTTNWYLGTYILYLDLATAHFQRMMPEEWTTIIHSLSKMFNLDSQLVL
EAY
~~~

–Sequence `S2`:

~~~
>gi|114341586|gb|ABI66866.1|
GRLQEFGVGEVTRESLRSMQAELPPVLEEALEVFYNTLSRAPEVDVLFRNDDHRAHAKRHQIKHWQRILS
GEYDTAYFDNVRRIGEVHFEIGLEPRYYVAGYAGIASSLVRGVIQSGQRNRVSRTKQFEATAAKVDALIR
AVFLDMELALSTY
~~~

–ProtT5-score end-gap-free alignment, which is identical with the reference MSA induced alignment:

~~~
S1 : 1 EQINYIGITDFDVDLLHSKETQFRAVVDMLVDELYEQITAQPELHRIILQHSTVERLKET 60
             G        L S       V        Y      PE              K
S2 : 1 GRLQEFGVGEVTRESLRSMQAELPPVLEEALEVFYNTLSRAPEVDVLFRNDDHRAHAKRH 60
S1 : 61 QRWYFLSMASGVINEPFIEKRLHIGKVHSRIGLTTNWYLGTYILYLDLATAHFQRMM 117
        Q        SG            IG VH  IGL        Y  Y
S2 : 61 QIKHWQRILSGEYDTAYFDNVRRIGEVHFEIGLEPRYYVAGYAGIASSLVRGVIQSGQRN 120
S1 : 118 -------PEEWTTIIHSLSKMFNLDSQLVLEAY 143
                 E        L     LD  L L  Y
S2 : 121 RVSRTKQFEATAAKVDALIRAVFLDMELALSTY 153
~~~

– BLOSUM45 matrix end-gap-free alignment:

~~~
S1 : 1 -EQINYIGITDFDVDLLHSKETQFRAVVDMLVDELYEQITAQPEL--------HRIILQH 51
        ++    G+ +   + L S + ++  V++   ++ +Y ++  PE+        HR H
S2 : 1 G-RLQEFGVGEVTRESLRSMQAELPPVLEEALEVFYNTLSRAPEVDVLFRNDDHRA---H 56
S1 : 52 STVERLKETQRWYFLSMASGVINEPFIEKRLHIGKVHSRIGLTTNWYLGTY------ILY 105
        +   ++K  QR     + SG  + + + IG+VH IGL +Y+ Y ++
S2 : 57 AKRHQIKHWQR-----ILSGEYDTAYFDNVRRIGEVHFEIGLEPRYYVAGYAGIASSLVR 111
S1 : 106 LDLATAHFQRM----MPEEWTTIIHSLSKMFNLDSQLVLEAY 143
           +  +  +  R+  E +  +L  +  LD  + L  L  Y
S2 : 112 GVIQSGQRNRVSRTKQFEATAAKVDALIRAVFLDMELALSTY 153
~~~

It appears that the BLOSUM45 matrix often introduces false gaps to achieve a higher number of matches. That is, it blindly works to achieve a maximum score while the ProtT5-score avoids this trap, due to its contextual nature. For example, note in the first example that BLOSUM45 removes one match to instead introduce six additional ones. Remarkably, the ProtT5-score finds the true alignment, in spite of the fact that it contains in the middle 21-18 region only one match instead of six. The ProtT5-score apears to be taking more information from the properties of the amino acids than are encoded in the matrix or capable of being encoded in the matrix without context.

### 3.3 Comparison procedure

We compare the *E*-scores against the BLOSUM matrices on the 49 MSAs introduced above as follows. For each MSA, we pick 300 randomly selected protein pairs (if the total number of pairs is less than 300, then we pick all pairs), compute pairwise alignment of each pair using either the *E*-score or BLO-SUM matrix, according to the Needleman-Wunsch dynamic programming global alignment algorithm [23], but without any penalties for gaps at the ends; we call this *end-gap-free* alignment. This is the most suitable alignment type, as we expect these sequences to align globally, however, using end-gap penalties could artificially alter the alignments.

For BLOSUM matrices we used the NCBI BLAST default values: − 11 for gap opening and − 1 for gap extension. For *E*-scores we used experimentally set gap penalties:. − 25 for gap opening and − 0.01 for gap extension. Experiments indicate that the algorithm is robust to penalty changes.

Each pairwise alignment is then evaluated against the reference alignment from CDD by computing the distance to the pairwise alignment induced by the reference MSA for the two sequences. The distance is computed using all five distances introduce above: d_cc_, d_d_, d_pos_, d_seq_, and d_ssp_.

The comparison for all *E*-scores and all BLOSUM matrices on all data is very time consuming, therefore, we need to choose the best *E*-score and the best BLOSUM matrix. For that purpose, we have selected six representative MSAs of the above 49; see the first column of Supplementary Tables 3-6. We have first compared all six *E*-scores on these six MSAs; the results are shown in Supplementary Table 3. The two top performers are the ProtT5-score and the ESM2-score. They are compared head-to-head in Supplementary Table 4, using the same six MSAs, but considering also the Wilcoxon signed-rank test P-values, with a threshold of 0.01, to check the validity of the findings. The result is that ProtT5 is the best *E*-score.

Similarly, to find the best BLOSUM matrix for our purposes, we performed the same procedure. All five candidates, BLOSUM45, BLOSUM50, BLOSUM62, BLOSUM80, and BLOSUM90 are compared in Supplementary Table 5, and the top two, BLOSUM45 and BLOSUM62, in Supplementary Table 6. BLOSUM45 is the best scoring matrix for our comparison.

### 3.4 ProtT5-score vs BLOSUM45 matrix

In this section we compare head-to-head the ProtT5-score against the BLOSUM45 matrix, the top performers, as decided by the previous section. We have compared the two, as described above, on all data; see Supplementary Table 1, where the Wilcoxon test P-values larger than .01 are indicated in red. We have removed all tests with P-values higher than .01 and present the comparison for the d_cc_, d_d_ and d_pos_ distances in Table 2. The d_cc_ distance presents the most important evaluation, which, together with d_d_ and d_pos_ gives a comprehensive evaluation of the similarity between the pairwise alignments and true ones, as induced by MSAs. The results for the d_ssp_ and d_seq_ distances are similar to those for the d_pos_ distance; see Supplementary Table 1.

**Table 2:** ProtT5-score against BLOSUM45 matrix: comparison using 38 MSAs, for which the Wilcoxon test P-values are smaller than .01. The best values are shown in boldface. We show only the distances d_cc_, d_d_, and d_pos_; the comparisons using d_ssp_ and d_seq_ are similar to the one for d_pos_.

| MSA | $d_{cc}$ | | | $d_d$ | | | $d_{pos}$ | | |
| --- | --- | --- | --- | --- | --- | --- | --- | --- | --- |
| Conserved domain | ProtT5 | BLOSUM45 | P-value | ProtT5 | BLOSUM45 | P-value | ProtT5 | BLOSUM45 | P-value |
| *Bbox2.TRIM37-C-VIII* | 0.00433 | **0.00339** | 2.4E-05 | 0.00133 | **0.00101** | 1.9E-07 | 0.05080 | **0.04291** | 4.9E-03 |
| *CBD-like* | **0.01111** | 0.02109 | 1.5E-26 | **0.00398** | 0.01075 | 4.4E-34 | **0.15886** | 0.23935 | 3.1E-32 |
| *Bbox2* | **0.01164** | 0.02593 | 2.9E-15 | **0.00297** | 0.00693 | 6.9E-14 | **0.13424** | 0.19607 | 3.5E-15 |
| *ChtBD1* | **0.01246** | 0.02158 | 2.5E-14 | **0.00319** | 0.00803 | 6.3E-16 | **0.15759** | 0.21879 | 3.6E-15 |
| *Bbox1_CYLD* | **0.01368** | 0.02840 | 1.8E-25 | **0.00451** | 0.00726 | 3.1E-09 | **0.15525** | 0.22111 | 1.4E-21 |
| *KAZAL_FS* | **0.01447** | 0.03913 | 1.1E-31 | **0.00345** | 0.01395 | 1.3E-31 | **0.14937** | 0.26945 | 1.6E-28 |
| *bHLH_SF* | **0.00718** | 0.02787 | 5.9E-25 | **0.00284** | 0.01238 | 3.8E-24 | **0.10214** | 0.21861 | 8.9E-23 |
| *CD_CSD* | **0.00949** | 0.02383 | 4.8E-25 | **0.00166** | 0.00617 | 7.8E-34 | **0.11058** | 0.20413 | 1.1E-29 |
| *C1* | **0.01392** | 0.02262 | 6.2E-20 | **0.00395** | 0.00826 | 3.6E-15 | **0.16602** | 0.22826 | 5.8E-16 |
| *TrHb* | **0.00453** | 0.04156 | 8.9E-05 | **0.00100** | 0.00768 | 1.4E-04 | **0.09407** | 0.34595 | 8.9E-05 |
| *Hb* | **0.00149** | 0.00249 | 5.1E-06 | **0.00020** | 0.00069 | 9.4E-09 | **0.04412** | 0.07279 | 4.6E-08 |
| *Hb-beta-like* | **0.00112** | 0.00200 | 2.5E-09 | **0.00018** | 0.00054 | 8.3E-13 | **0.02340** | 0.03875 | 1.2E-12 |
| *Hb-alpha-like* | **0.00068** | 0.00097 | 1.9E-03 | **0.00004** | 0.00023 | 1.4E-05 | **0.01533** | 0.02272 | 8.5E-04 |
| *GS_GGDEF_2* | **0.00184** | 0.01204 | 1.7E-45 | **0.00027** | 0.00337 | 5.5E-44 | **0.02721** | 0.16645 | 8.0E-48 |
| *Globin-like* | **0.03239** | 0.16054 | 6.3E-21 | **0.00655** | 0.11860 | 5.4E-21 | **0.37020** | 0.80690 | 6.0E-21 |
| *FHb-globin* | **0.00123** | 0.00221 | 1.8E-13 | **0.00015** | 0.00058 | 1.4E-28 | **0.02516** | 0.04282 | 4.6E-25 |
| *PFM_HFR-2-like* | **0.00627** | 0.00856 | 9.1E-05 | **0.00152** | 0.00194 | 7.5E-08 | **0.09867** | 0.12925 | 2.4E-14 |
| *SH2_STAT_family* | **0.01980** | 0.03637 | 3.9E-31 | **0.00584** | 0.01336 | 1.1E-21 | **0.20702** | 0.29659 | 2.0E-33 |
| *PBP-like* | **0.01190** | 0.03805 | 1.0E-43 | **0.00331** | 0.01568 | 3.8E-37 | **0.19009** | 0.34945 | 6.7E-45 |
| *SH2* | **0.03439** | 0.07021 | 2.3E-38 | **0.00929** | 0.03095 | 1.9E-35 | **0.30388** | 0.48456 | 3.9E-43 |
| *Globin_sensor* | **0.00456** | 0.02314 | 1.3E-47 | **0.00059** | 0.00749 | 8.8E-49 | **0.06854** | 0.25423 | 3.4E-48 |
| *PFM_monalysin-like* | **0.01356** | 0.01989 | 2.2E-19 | **0.00505** | 0.00563 | 1.4E-08 | **0.26374** | 0.30890 | 3.9E-18 |
| *Mb-like* | **0.00886** | 0.04507 | 3.2E-49 | **0.00225** | 0.01988 | 2.2E-47 | **0.16732** | 0.42281 | 2.1E-49 |
| *FYVE-like_SF* | **0.01218** | 0.04664 | 1.3E-43 | **0.00531** | 0.02462 | 4.2E-42 | **0.18193** | 0.32957 | 5.3E-44 |
| *PFM_aerolysin_family* | **0.04624** | 0.08305 | 7.7E-48 | **0.01082** | 0.03743 | 1.2E-44 | **0.64751** | 0.77877 | 5.4E-39 |
| *7tmA_photoreceptors_insect* | **0.00140** | 0.00245 | 2.9E-34 | **0.00023** | 0.00108 | 1.6E-40 | **0.04946** | 0.08400 | 5.3E-39 |
| *7tmA_Melanopsin-like* | **0.00088** | 0.00387 | 2.1E-10 | **0.00013** | 0.00184 | 1.9E-10 | **0.04305** | 0.11684 | 2.1E-10 |
| *ClyA_AhlB-like* | **0.00200** | 0.00398 | 9.5E-26 | **0.00019** | 0.00090 | 1.4E-38 | **0.03305** | 0.07839 | 1.3E-40 |
| *7tmA_Opsins_type2_animals* | **0.00200** | 0.00824 | 4.6E-50 | **0.00056** | 0.00372 | 1.1E-48 | **0.06925** | 0.23179 | 1.3E-50 |
| *7tm_GPCRs* | **0.02268** | 0.09884 | 4.6E-24 | **0.00533** | 0.07128 | 5.6E-24 | **0.53322** | 0.89102 | 6.3E-24 |
| *7tmA_Anaphylatoxin_R-like* | **0.00393** | 0.01085 | 1.6E-23 | **0.00119** | 0.00369 | 3.8E-22 | **0.10993** | 0.21479 | 6.7E-24 |
| *7tmA_Opioid_R-like* | **0.00204** | 0.00713 | 7.8E-19 | **0.00097** | 0.00341 | 2.7E-17 | **0.11747** | 0.19390 | 3.5E-18 |
| *ClyA-like* | **0.02331** | 0.06510 | 8.0E-48 | **0.01372** | 0.03956 | 3.9E-46 | **0.43093** | 0.67044 | 2.0E-49 |
| *FGGY* | **0.00458** | 0.01142 | 7.4E-49 | **0.00227** | 0.00490 | 9.1E-35 | **0.18887** | 0.35974 | 1.2E-49 |
| *FGGY_YpCarbK-like* | **0.00160** | 0.00486 | 3.8E-39 | **0.00022** | 0.00140 | 1.6E-45 | **0.06318** | 0.10790 | 1.0E-48 |
| *7tm_classA_rhodopsin-like* | **0.00605** | 0.02588 | 6.5E-51 | **0.00206** | 0.01231 | 1.2E-49 | **0.18837** | 0.44775 | 6.7E-51 |
| *NBD_sugar-kinase_HSP70_actin* | **0.03337** | 0.12052 | 6.4E-50 | **0.01808** | 0.10443 | 1.2E-48 | **0.66400** | 0.90307 | 1.3E-50 |
| *7tmA_amine_R-like* | **0.00475** | 0.02263 | 5.7E-50 | **0.00240** | 0.01630 | 8.1E-47 | **0.18416** | 0.35179 | 9.7E-51 |

**Table 3:** ProtAlbert-score against BLOSUM45 matrix: comparison using 6 MSAs. The best values are shown in boldface. We show only the distances d_cc_, d_d_, and d_pos_; the comparisons using d_ssp_ and d_seq_ are similar to the one for d_pos_.

| MSA | $d_{cc}$ | | | $d_d$ | | | $d_{pos}$ | | |
| --- | --- | --- | --- | --- | --- | --- | --- | --- | --- |
| Conserved domain | ProtAlbert | BLOSUM45 | P-value | ProtAlbert | BLOSUM45 | P-value | ProtAlbert | BLOSUM45 | P-value |
| *Hb-alpha-like* | **0.00064** | 0.00097 | 2.9E-05 | **0.00005** | 0.00023 | 2.4E-07 | **0.01538** | 0.02272 | 7.2E-06 |
| *Globin-like* | **0.04961** | 0.16054 | 1.5E-19 | **0.01144** | 0.11860 | 3.3E-19 | **0.54422** | 0.80690 | 6.1E-20 |
| *7tmA_photoreceptors_insect* | **0.00139** | 0.00245 | 1.3E-38 | **0.00018** | 0.00108 | 1.4E-43 | **0.04539** | 0.08400 | 1.1E-42 |
| *7tm_GPCRs* | **0.03310** | 0.09884 | 4.9E-24 | **0.00908** | 0.07128 | 9.6E-23 | **0.66807** | 0.89102 | 1.7E-23 |
| *FGGY_YpCarbK-like* | **0.00256** | 0.00486 | 7.3E-29 | **0.00038** | 0.00140 | 1.9E-44 | **0.07149** | 0.10790 | 2.9E-47 |
| *NBD_sugar-kinase_HSP70_actin* | **0.04928** | 0.12052 | 1.2E-48 | **0.03440** | 0.10443 | 3.6E-42 | **0.78501** | 0.90307 | 4.9E-47 |

ProtT5-score is better in all but one test on the shortest MSA. Regarding the overall number of better cases, ProtT5 has 73% of the cases, BLOSUM45 13%, while for the remaining 14% they were equal; see Supplementary Table 2. However, the distribution of these better cases, together with the performance on each individual case, are so that the ProtT5-score performs better in all but the first case on the average. For a better visualization of the comparison, we plotted in Figure 4, for each of the five distances, the ratio between the distance to the reference alignment from the BLOSUM45 alignment and from the ProtT5 alignment, respectively. That means higher than 1 is better in favour of ProtT5. It is very clear from the figure that ProtT5 has a much better similarity with the reference alignment, the advantage being most clear for the d_d_ distance, as expected.

**Figure 4.**
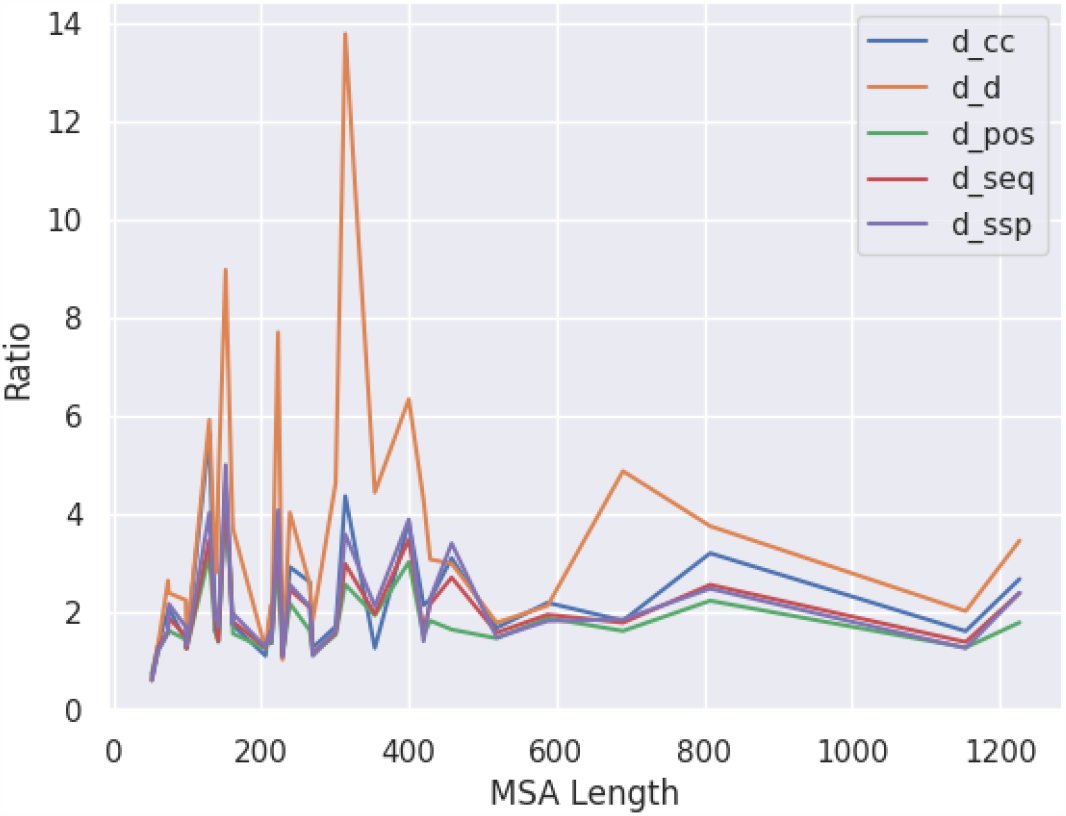
The ratio between the performance of BLOSUM45 and ProtT5-score in terms of distances to the reference from their alignments. Higher than 1 indicates ProtT5-score is better.

The comparison for the purely global alignment case (where end-gaps are considered) is presented in the Supplementary Table 7. The conclusions are similar to those presented here for the end-gap-free case.

### 3.5 ProtAlbert-score vs BLOSUM45 matrix

We have seen above that the best *E*-score, ProtT5, is much better than the best BLOSUM matrix, BLOSUM45. However, it turns out that the least powerful *E*-score, ProtAlbert (strictly speaking, ProtXLNet is the least powerful, but it gives errors sometimes, and cannot be tested properly) is still clearly outperforming the top BLOSUM45 matrix. We show only the comparison on the six selected alignments in Table 3. This indicates most clearly the superiority of the *E*-scores compared with BLOSUM matrices for scoring alignments.

## 4 Discussion and conclusion

We have proposed a new method for scoring sequence alignments, *E*-score, using the cosine similarity between the vectors associated using any word embedding algorithm, *E*. When using contextual embeddings, the *E*-score has the advantage of incorporating into the score associated with each position contextual information from the neighbouring positions in the sequence.

We have thoroughly tested the new *E*-score in the context of protein alignments, by comparing the top scoring function, ProtT5-score, against the BLOSUM45 matrix on a large set including a wide variety of reference multiple sequence alignments from NCBI’s Conserved Domain Database. The pairwise alignments produced by using the ProtT5-score are much closer to the true alignments, as induced by the reference multiple sequence alignments, compared to BLOSUM45. Furthermore, even the least powerful ProtAlbert-score still clearly outperforms the top BLOSUM45 matrix.

While we have tested the new *E*-score in the context of biological sequences, particularly proteins, the new method can be applied to any sequences where computation of embedding vectors is possible, which include practically all existing textual data, the largest type of data available. All Large Language Models (LLM) have the possibility to produce word embedding vectors for any input text. Considering the importance and implications of text similarity in general, the potential for applications of our new method appears to be virtually boundless.

The language models are expected to improve at a rapid pace in the future, with expected corresponding improvements for those involving biological sequences. The ideas presented apply directly to any new model, meaning that the *E*-score will improve as well.

Specialized models, obtained usually by fine tuning the general ones, or adapting them to specialized tasks, are expected to provide improved performance. Regarding the case of biological sequences, where alignments play a particularly vital role, the possibility exists for training, or fine tuning, models for highly specialized tasks, such as specific types of proteins.

Finally, aligning nucleic acid sequences – DNA, RNA – though simpler than the case of proteins, is a whole area where *E*-scores can be applied as well.

## 5 Availability

The program to compute alignments based on various *E*-scores is available as a web server at e-score.csd.uwo.ca. The source code is freely available for download from github.com/lucian-ilie/E-score.

## 6 Author contributions statement

S.A. implemented the alignment programs, downloaded and installed the embedding models, downloaded and processed the data, performed all testing, and built the GitHub page and web server. G.B.G. proposed the Conserved Domain Database, selected all reference MSAs used for testing, identified the six representative ones used for pre-testing, and wrote the description of the data. S.I. introduced the new closest context distance and contributed to finding the gap penalties. L.I. proposed the problem, defined the *E*-score, proposed the evaluation using pairwise alignments induced by MSAs, introduced the new relative displacement distance, supervised the implementation and testing, and wrote the manuscript. All authors read and approved the final version of the manuscript.

## 7. Acknowledgments

SeyedMohsen Hosseini provided assistance with building the web server and installing some embedding models. All computations were performed on Compute Canada servers.

## 9 Supplementary data

A supplementary file containing Supplementary Tables 1-8 is available online.

## Footnotes

1 The two BLAST papers [2, 3] combined exceed 190,000 citations on Google Scholar as of Aug. 2023.

