## Supplementary Tables for "Scoring alignments by embedding vector similarity"

– Supplementary material –

Sepehr Ashrafzadeh, G. Brian Golding, Silvana Ilie, Lucian Ilie*

November 16, 2023

#### 1 End-gap-free alignment tests

Supplementary Table 1: End-gap-free alignments: ProtT5-score vs BLOSUM45 matrix, average distances for all five distances and all testing MSAs. Best results are shown in boldface. Wilcoxon test P-values higher than .01 are shown in red.

| MSA | | | | | d_cc | | | d_d | | | d_pos | | | d_seq | | | d_ssp | | |
|---|---|---|---|---|---|---|---|---|---|---|---|---|---|---|---|---|---|---|---|
| Conserved domain | Source | Proteins | Length | Samples | ProtT5 | BLOSUM45 | P-value | ProtT5 | BLOSUM45 | P-value | ProtT5 | BLOSUM45 | P-value | ProtT5 | BLOSUM45 | P-value | ProtT5 | BLOSUM45 | P-value |
| Bbox2_MID2_C-I | cd19823 | 8 | 40 | 21 | 0.000000 | 0.000000 | - | 0.000000 | 0.000000 | - | 0.000000 | 0.000000 | - | 0.000000 | 0.000000 | - | 0.000000 | 0.000000 | - |
| Bbox2_TRIM42_C-III | cd19782 | 9 | 40 | 28 | **0.003005** | 0.007226 | 1.23E-01 | 0.000174 | **0.001647** | 1.17E-02 | **0.016727** | 0.045626 | 1.15E-02 | **0.016727** | 0.045626 | 1.15E-02 | **0.020341** | 0.030863 | 3.98E-01 |
| Bbox2_MID | cd19758 | 10 | 40 | 36 | **0.001193** | 0.001380 | 7.20E-02 | **0.000361** | 0.000577 | 8.98E-03 | **0.033052** | 0.040788 | 3.79E-02 | **0.033052** | 0.040788 | 3.79E-02 | 0.044817 | **0.036704** | 6.36E-01 |
| Bbox2_MID1_C-I | cd19822 | 9 | 47 | 28 | 0.000000 | 0.000000 | - | 0.000000 | 0.000000 | - | 0.000000 | 0.000000 | - | 0.000000 | 0.000000 | - | 0.000000 | 0.000000 | - |
| Bbox_SF | cd00021 | 6 | 48 | 10 | **0.023063** | 0.048396 | 1.10E-01 | **0.011057** | 0.017582 | 6.78E-01 | **0.236113** | 0.334374 | 1.39E-01 | **0.234950** | 0.319753 | 1.73E-01 | **0.241883** | 0.324529 | 3.74E-01 |
| DEFL_defensin-like | cd21806 | 78 | 51 | 300 | 0.010198 | **0.009544** | 5.64E-04 | **0.003007** | 0.003981 | 4.04E-01 | **0.119352** | 0.119963 | 5.47E-01 | **0.104636** | 0.105939 | 6.51E-01 | **0.098153** | 0.103229 | 6.41E-01 |
| Bbox2_TRIM37_C-VIII | cd19779 | 25 | 52 | 276 | 0.004326 | **0.003388** | 2.39E-05 | 0.001326 | **0.001015** | 1.92E-07 | 0.050800 | **0.042909** | 4.87E-03 | 0.044965 | **0.035128** | 1.36E-04 | 0.037020 | **0.022129** | 3.22E-13 |
| Bbox2_TRIM9-like | cd19764 | 17 | 53 | 120 | **0.006689** | 0.011801 | 1.48E-02 | **0.001876** | 0.003876 | 6.02E-05 | **0.066398** | 0.099693 | 2.24E-06 | **0.052880** | 0.088069 | 1.27E-06 | **0.049004** | 0.078140 | 4.13E-03 |
| CBD_like | cd12204 | 41 | 61 | 300 | **0.011107** | 0.021086 | 1.46E-26 | **0.003980** | 0.010755 | 4.38E-34 | **0.158857** | 0.239354 | 3.10E-32 | **0.136414** | 0.219661 | 7.40E-33 | **0.162183** | 0.220194 | 6.95E-17 |
| Bbox2 | cd19756 | 127 | 65 | 300 | **0.011640** | 0.025928 | 2.87E-15 | **0.002966** | 0.006933 | 6.86E-14 | **0.134239** | 0.196068 | 3.54E-15 | **0.127539** | 0.188995 | 6.74E-15 | **0.143830** | 0.203194 | 3.55E-07 |
| ChtBD1 | cd00035 | 32 | 67 | 300 | **0.012463** | 0.021581 | 2.51E-14 | **0.003191** | 0.008026 | 6.35E-16 | **0.157591** | 0.218785 | 3.59E-15 | **0.128995** | 0.187655 | 5.84E-15 | **0.124513** | 0.185850 | 2.51E-12 |
| Bbox1_CYLD | cd19816 | 27 | 68 | 325 | **0.013685** | 0.028400 | 1.85E-25 | **0.004505** | 0.007261 | 3.06E-09 | **0.155255** | 0.221111 | 1.36E-21 | **0.146511** | 0.212084 | 5.98E-21 | **0.165157** | 0.237162 | 2.90E-16 |
| KAZAL_FS | cd00104 | 273 | 74 | 300 | **0.014470** | 0.039131 | 1.06E-31 | **0.003451** | 0.013951 | 1.33E-31 | **0.149365** | 0.269448 | 1.61E-28 | **0.140465** | 0.257798 | 3.75E-28 | **0.163934** | 0.285377 | 1.47E-22 |
| bHLH_SF | cd00083 | 79 | 75 | 300 | **0.007183** | 0.027871 | 5.85E-25 | **0.002838** | 0.012381 | 3.77E-24 | **0.102140** | 0.218614 | 8.85E-23 | **0.079882** | 0.206034 | 3.25E-24 | **0.079658** | 0.228492 | 2.62E-23 |
| CD_CSD | cd00024 | 522 | 98 | 300 | **0.009485** | 0.023830 | 4.80E-25 | **0.001658** | 0.006172 | 7.85E-34 | **0.110584** | 0.204128 | 1.06E-29 | **0.100455** | 0.196800 | 9.41E-31 | **0.105495** | 0.229457 | 7.07E-30 |
| C1 | cd00029 | 281 | 99 | 300 | **0.013915** | 0.022624 | 6.22E-20 | **0.003952** | 0.008256 | 3.58E-15 | **0.166022** | 0.228262 | 5.84E-16 | **0.157327** | 0.217656 | 1.14E-14 | **0.175744** | 0.246968 | 7.70E-13 |
| TrHb | cd14756 | 8 | 130 | 21 | **0.004535** | 0.041555 | 8.86E-05 | **0.001002** | 0.007677 | 1.40E-04 | **0.094067** | 0.345955 | 8.86E-05 | **0.083484** | 0.338490 | 8.84E-05 | **0.092226** | 0.403291 | 8.86E-05 |
| Hb | cd14765 | 15 | 138 | 91 | **0.001490** | 0.002487 | 5.10E-06 | **0.000204** | 0.000687 | 9.39E-09 | **0.044115** | 0.072793 | 4.59E-08 | **0.034136** | 0.066287 | 5.17E-10 | **0.041652** | 0.079085 | 5.05E-09 |
| SH2_STAT5 | cd10376 | 5 | 140 | 6 | **0.001355** | 0.002005 | 4.58E-01 | **0.000121** | 0.000146 | 4.58E-01 | 0.020036 | **0.017497** | 4.58E-01 | 0.018824 | **0.016285** | 4.58E-01 | 0.015888 | **0.011003** | 4.58E-01 |
| Hb-beta-like | cd08925 | 27 | 140 | 300 | **0.001115** | 0.001996 | 2.49E-09 | **0.000179** | 0.000536 | 8.30E-13 | **0.023404** | 0.038748 | 1.16E-12 | **0.021218** | 0.036147 | 7.33E-12 | **0.020491** | 0.041377 | 1.78E-11 |
| Hb-alpha-like | cd08927 | 39 | 142 | 300 | **0.000678** | 0.000975 | 1.87E-03 | **0.000044** | 0.000228 | 1.40E-05 | **0.015327** | 0.022717 | 8.54E-04 | **0.014910** | 0.022455 | 6.91E-04 | **0.015378** | 0.027156 | 2.36E-04 |
| SH2_STAT5a | cd10421 | 10 | 145 | 36 | **0.001055** | 0.006663 | 4.57E-02 | **0.000129** | 0.000639 | 4.53E-02 | **0.014668** | 0.032758 | 4.53E-02 | **0.011324** | 0.030027 | 4.53E-02 | **0.007680** | 0.034564 | 4.53E-02 |
| Mb | cd08926 | 9 | 149 | 28 | 0.000681 | **0.000673** | 8.33E-01 | **0.000049** | 0.000086 | 4.35E-04 | **0.014150** | 0.021647 | 3.82E-04 | **0.013665** | 0.021647 | 3.83E-04 | **0.010430** | 0.018105 | 3.44E-02 |
| GS_GGDEF_2 | cd14759 | 26 | 152 | 300 | **0.001836** | 0.012038 | 1.71E-45 | **0.000270** | 0.003370 | 5.51E-44 | **0.027214** | 0.166451 | 8.01E-48 | **0.026258** | 0.165448 | 1.16E-47 | **0.034206** | 0.209150 | 6.81E-42 |
| Globin-like | cd01067 | 17 | 161 | 120 | **0.032387** | 0.160539 | 6.33E-21 | **0.006550** | 0.118599 | 5.43E-21 | **0.370198** | 0.806900 | 6.02E-21 | **0.349277** | 0.775463 | 5.30E-21 | **0.410540** | 0.823240 | 4.91E-21 |
| FHb-globin | cd08922 | 47 | 162 | 300 | **0.001234** | 0.002214 | 1.81E-13 | **0.000146** | 0.000584 | 1.43E-28 | **0.025158** | 0.042822 | 4.57E-25 | **0.019062** | 0.038008 | 4.92E-26 | **0.018162** | 0.037874 | 1.16E-22 |
| PFM_HFR-2-like | cd20216 | 90 | 174 | 300 | **0.006271** | 0.008556 | 9.07E-05 | **0.001516** | 0.001944 | 7.48E-08 | **0.098671** | 0.129252 | 2.38E-14 | **0.097780** | 0.127898 | 5.27E-14 | **0.138235** | 0.175490 | 9.29E-12 |
| SH2_STAT_family | cd09919 | 67 | 206 | 300 | **0.019796** | 0.036365 | 3.91E-31 | **0.005837** | 0.013364 | 1.12E-21 | **0.207023** | 0.296589 | 2.03E-33 | **0.182054** | 0.274387 | 1.93E-33 | **0.211894** | 0.304906 | 4.73E-29 |
| PBP-like | cd08919 | 31 | 213 | 300 | **0.011904** | 0.038051 | 1.02E-43 | **0.003310** | 0.015676 | 3.77E-37 | **0.190088** | 0.349447 | 6.69E-45 | **0.177615** | 0.336765 | 3.42E-45 | **0.216576** | 0.393467 | 4.63E-43 |
| SH2 | cd00173 | 352 | 214 | 300 | **0.034392** | 0.070210 | 2.32E-38 | **0.009287** | 0.030954 | 1.87E-35 | **0.303880** | 0.484562 | 3.94E-43 | **0.279447** | 0.460347 | 6.69E-43 | **0.333277** | 0.525209 | 6.60E-39 |
| Globin_sensor | cd01068 | 193 | 223 | 300 | **0.004564** | 0.023144 | 1.30E-47 | **0.000586** | 0.007488 | 8.78E-49 | **0.068540** | 0.254229 | 3.37E-48 | **0.052203** | 0.244672 | 2.05E-48 | **0.062197** | 0.294697 | 1.93E-47 |
| PFM_monalysin-like | cd17904 | 31 | 229 | 300 | **0.013563** | 0.019888 | 2.23E-19 | **0.005046** | 0.005629 | 1.43E-08 | **0.263737** | 0.308905 | 3.85E-18 | **0.261511** | 0.306512 | 9.59E-18 | **0.346435** | 0.390074 | 8.77E-15 |
| Mb-like | cd01040 | 384 | 239 | 300 | **0.008856** | 0.045075 | 3.23E-49 | **0.002254** | 0.019879 | 2.17E-47 | **0.167324** | 0.422814 | 2.10E-49 | **0.142339** | 0.408479 | 2.50E-49 | **0.164794** | 0.470599 | 6.41E-49 |
| FYVE_like_SF | cd00065 | 392 | 266 | 300 | **0.012183** | 0.046637 | 1.30E-43 | **0.005308** | 0.024618 | 4.23E-42 | **0.181928** | 0.329575 | 5.34E-44 | **0.108460** | 0.262068 | 1.16E-45 | **0.124533** | 0.294140 | 3.34E-42 |
| PFM_aerolysin_family | cd10140 | 65 | 270 | 300 | **0.046240** | 0.083055 | 7.71E-48 | **0.010819** | 0.037435 | 1.20E-44 | **0.647508** | 0.778774 | 5.43E-39 | **0.613236** | 0.744310 | 1.82E-38 | **0.723727** | 0.828257 | 1.00E-33 |
| 7tmA_photoreceptors_insect | cd15079 | 35 | 301 | 300 | **0.001399** | 0.002451 | 2.89E-34 | **0.000228** | 0.001084 | 1.58E-40 | **0.049456** | 0.084005 | 5.32E-39 | **0.047164** | 0.082066 | 2.64E-39 | **0.065482** | 0.114693 | 3.38E-36 |
| 7tmA_Melanopsin-like | cd15083 | 12 | 314 | 55 | **0.000879** | 0.003872 | 2.15E-10 | **0.000131** | 0.001837 | 1.92E-10 | **0.043049** | 0.116844 | 2.15E-10 | **0.034671** | 0.110080 | 1.92E-10 | **0.038653** | 0.144452 | 2.40E-10 |
| ClyA_AhlB-like | cd22652 | 39 | 354 | 300 | **0.001996** | 0.003982 | 9.53E-26 | **0.000187** | 0.000903 | 1.43E-38 | **0.033045** | 0.078389 | 1.32E-40 | **0.031856** | 0.076990 | 5.07E-40 | **0.042178** | 0.105673 | 2.45E-32 |
| 7tmA_Opsins_type2_animals | cd14969 | 71 | 400 | 300 | **0.001996** | 0.008244 | 4.64E-50 | **0.000562** | 0.003724 | 1.09E-48 | **0.069253** | 0.231791 | 1.33E-50 | **0.057024** | 0.221317 | 1.33E-50 | **0.067398** | 0.286427 | 2.15E-50 |
| 7tm_GPCRs | cd14964 | 18 | 420 | 136 | **0.022676** | 0.098840 | 4.60E-24 | **0.005327** | 0.071275 | 5.61E-24 | **0.533216** | 0.891019 | 6.27E-24 | **0.480151** | 0.849745 | 5.74E-24 | **0.609748** | 0.912477 | 8.17E-24 |
| 7tmA_Anaphylatoxin_R-like | cd14974 | 18 | 429 | 136 | **0.003931** | 0.010849 | 1.56E-23 | **0.001186** | 0.003688 | 3.79E-22 | **0.109928** | 0.214792 | 6.71E-24 | **0.080396** | 0.188687 | 6.71E-24 | **0.102004** | 0.247353 | 7.51E-24 |
| 7tmA_Opioid_R-like | cd14970 | 16 | 458 | 105 | **0.002042** | 0.007125 | 7.78E-19 | **0.000973** | 0.003406 | 2.67E-17 | **0.117470** | 0.193901 | 3.50E-18 | **0.047128** | 0.130043 | 1.25E-18 | **0.048755** | 0.167504 | 8.78E-19 |
| ClyA-like | cd21116 | 118 | 519 | 300 | **0.023307** | 0.065095 | 8.05E-48 | **0.013722** | 0.039557 | 3.92E-46 | **0.430935** | 0.670442 | 2.03E-49 | **0.377331** | 0.646181 | 1.19E-49 | **0.463617** | 0.711213 | 9.36E-49 |
| FGGY_RBK_like | cd07768 | 6 | 537 | 10 | **0.005103** | 0.008179 | 1.95E-03 | **0.001284** | 0.003311 | 1.95E-02 | **0.227322** | 0.315757 | 1.95E-03 | **0.206354** | 0.303679 | 1.95E-03 | **0.282124** | 0.381714 | 1.95E-03 |
| FGGY | cd00366 | 31 | 588 | 300 | **0.004575** | 0.011417 | 7.38E-49 | **0.002272** | 0.004898 | 9.09E-35 | **0.188870** | 0.359736 | 1.24E-49 | **0.178342** | 0.352315 | 1.24E-49 | **0.238063** | 0.435521 | 1.49E-49 |
| FGGY_YpCarbK_like | cd07782 | 35 | 690 | 300 | **0.001603** | 0.004856 | 3.77E-39 | **0.000224** | 0.001399 | 1.56E-45 | **0.063175** | 0.107904 | 1.01E-48 | **0.050921** | 0.097829 | 1.66E-48 | **0.062732** | 0.121254 | 1.00E-47 |
| 7tm_classA_rhodopsin-like | cd00637 | 405 | 808 | 300 | **0.006052** | 0.025884 | 6.46E-51 | **0.002059** | 0.012306 | 1.20E-49 | **0.188373** | 0.447748 | 6.66E-51 | **0.154920** | 0.426119 | 6.59E-51 | **0.201482** | 0.523437 | 6.86E-51 |
| NBD_sugar-kinase_HSP70_actin | cd00012 | 125 | 1154 | 300 | **0.033371** | 0.120524 | 6.40E-50 | **0.018078** | 0.104434 | 1.22E-48 | **0.664004** | 0.903067 | 1.29E-50 | **0.565443** | 0.825004 | 1.29E-50 | **0.688703** | 0.913651 | 1.29E-50 |
| 7tmA_amine_R-like | cd14967 | 78 | 1227 | 300 | **0.004752** | 0.022627 | 5.69E-50 | **0.002395** | 0.016302 | 8.13E-47 | **0.184162** | 0.351788 | 9.71E-51 | **0.106436** | 0.280950 | 1.06E-50 | **0.137242** | 0.352901 | 7.82E-51 |

Supplementary Table 2: End-gap-free alignments: ProtT5-score vs BLOSUM45 matrix, better cases.

| MSA | | | | | d_cc | | | d_d | | | d_pos | | | d_seq | | | d_ssp | | |
|---|---|---|---|---|---|---|---|---|---|---|---|---|---|---|---|---|---|---|---|
| **Conserved domain** | **Source** | **Proteins** | **Length** | **Samples** | **ProtT5** | **BLOSUM45** | Equal | **ProtT5** | **BLOSUM45** | Equal | **ProtT5** | **BLOSUM45** | Equal | **ProtT5** | **BLOSUM45** | Equal | **ProtT5** | **BLOSUM45** | Equal |
| Bbox2_MID2_C-I | cd19823 | 8 | 40 | 21 | 0 | 0 | 21 | 0 | 0 | 21 | 0 | 0 | 21 | 0 | 0 | 21 | 0 | 0 | 21 |
| Bbox2_TRIM42_C-III | cd19782 | 9 | 40 | 28 | **6** | 2 | 20 | **8** | 0 | 20 | **8** | 0 | 20 | **8** | 0 | 20 | **5** | 2 | 21 |
| Bbox2_MID | cd19758 | 10 | 40 | 36 | **11** | 3 | 22 | **11** | 1 | 24 | **10** | 2 | 24 | **10** | 2 | 24 | **9** | 5 | 22 |
| Bbox2_MID1_C-I | cd19822 | 9 | 47 | 28 | 0 | 0 | 28 | 0 | 0 | 28 | 0 | 0 | 28 | 0 | 0 | 28 | 0 | 0 | 28 |
| Bbox_SF | cd00021 | 6 | 48 | 10 | **6** | 3 | 1 | **5** | 4 | 1 | **6** | 3 | 1 | **6** | 3 | 1 | **6** | 3 | 1 |
| DEFL_defensin-like | cd21806 | 78 | 51 | 300 | 81 | **140** | 79 | 88 | **127** | 85 | 84 | **115** | 101 | 85 | **118** | 97 | 85 | **119** | 96 |
| Bbox2_TRIM37_C-VIII | cd19779 | 25 | 52 | 276 | 58 | **77** | 141 | 45 | **83** | 148 | 45 | **62** | 169 | 46 | **76** | 154 | 9 | **80** | 187 |
| Bbox2_TRIM9-like | cd19764 | 17 | 53 | 120 | **60** | 25 | 35 | **64** | 17 | 39 | **67** | 15 | 38 | **67** | 17 | 36 | **43** | 17 | 60 |
| CBD_like | cd12204 | 41 | 61 | 300 | **205** | 54 | 41 | **210** | 42 | 48 | **209** | 37 | 54 | **209** | 37 | 54 | **170** | 62 | 68 |
| Bbox2 | cd19756 | 127 | 65 | 300 | **179** | 79 | 42 | **169** | 74 | 57 | **167** | 64 | 69 | **168** | 67 | 65 | **148** | 87 | 65 |
| ChtBD1 | cd00035 | 32 | 67 | 300 | **170** | 83 | 47 | **164** | 80 | 56 | **159** | 73 | 68 | **157** | 70 | 73 | **149** | 81 | 70 |
| Bbox1_CYLD | cd19816 | 27 | 68 | 325 | **217** | 68 | 40 | **178** | 98 | 49 | **195** | 67 | 63 | **198** | 69 | 58 | **186** | 80 | 59 |
| KAZAL_FS | cd00104 | 273 | 74 | 300 | **209** | 41 | 50 | **201** | 45 | 54 | **199** | 40 | 61 | **200** | 43 | 57 | **179** | 68 | 53 |
| bHLH_SF | cd00083 | 79 | 75 | 300 | **174** | 55 | 71 | **173** | 52 | 75 | **173** | 48 | 79 | **172** | 48 | 80 | **170** | 53 | 77 |
| CD_CSD | cd00024 | 522 | 98 | 300 | **211** | 52 | 37 | **222** | 34 | 44 | **211** | 38 | 51 | **216** | 38 | 46 | **215** | 40 | 45 |
| C1 | cd00029 | 281 | 99 | 300 | **201** | 76 | 23 | **180** | 91 | 29 | **181** | 89 | 30 | **180** | 90 | 30 | **182** | 90 | 28 |
| TrHb | cd14756 | 8 | 130 | 21 | **20** | 0 | 1 | **19** | 1 | 1 | **20** | 0 | 1 | **20** | 0 | 1 | **20** | 0 | 1 |
| Hb | cd14765 | 15 | 138 | 91 | **51** | 16 | 24 | **52** | 15 | 24 | **51** | 16 | 24 | **51** | 13 | 27 | **51** | 13 | 27 |
| SH2_STAT5 | cd10376 | 5 | 140 | 6 | 2 | 2 | 2 | 2 | 2 | 2 | 2 | 2 | 2 | 2 | 2 | 2 | 2 | 2 | 2 |
| Hb-beta-like | cd08925 | 27 | 140 | 300 | **124** | 52 | 124 | **119** | 55 | 126 | **124** | 49 | 127 | **124** | 50 | 126 | **115** | 48 | 137 |
| Hb-alpha-like | cd08927 | 39 | 142 | 300 | **68** | 59 | 173 | **72** | 53 | 175 | **71** | 53 | 176 | **71** | 53 | 176 | **72** | 53 | 175 |
| SH2_STAT5a | cd10421 | 10 | 145 | 36 | **8** | 6 | 22 | **8** | 6 | 22 | **8** | 6 | 22 | **8** | 6 | 22 | **8** | 6 | 22 |
| Mb | cd08926 | 9 | 149 | 28 | **13** | 9 | 6 | **16** | 0 | 12 | **16** | 4 | 8 | **16** | 4 | 8 | **12** | 6 | 10 |
| GS_GGDEF_2 | cd14759 | 26 | 152 | 300 | **276** | 8 | 16 | **274** | 10 | 16 | **279** | 5 | 16 | **279** | 5 | 16 | **244** | 8 | 48 |
| Globin-like | cd01067 | 17 | 161 | 120 | **116** | 2 | 2 | **116** | 2 | 2 | **116** | 2 | 2 | **116** | 2 | 2 | **116** | 2 | 2 |
| FHb-globin | cd08922 | 47 | 162 | 300 | **163** | 63 | 74 | **178** | 35 | 87 | **176** | 34 | 90 | **177** | 34 | 89 | **165** | 37 | 98 |
| PFM_HFR-2-like | cd20216 | 90 | 174 | 300 | **140** | 96 | 64 | **150** | 85 | 65 | **163** | 67 | 70 | **161** | 69 | 70 | **157** | 78 | 65 |
| SH2_STAT_family | cd09919 | 67 | 206 | 300 | **223** | 58 | 19 | **200** | 81 | 19 | **223** | 49 | 28 | **225** | 50 | 25 | **213** | 66 | 21 |
| PBP-like | cd08919 | 31 | 213 | 300 | **266** | 23 | 11 | **253** | 36 | 11 | **259** | 22 | 19 | **264** | 21 | 15 | **252** | 33 | 15 |
| SH2 | cd00173 | 352 | 214 | 300 | **257** | 43 | 0 | **258** | 42 | 0 | **262** | 27 | 11 | **265** | 29 | 6 | **257** | 40 | 3 |
| Globin_sensor | cd01068 | 193 | 223 | 300 | **275** | 16 | 9 | **280** | 10 | 10 | **279** | 8 | 13 | **280** | 8 | 12 | **272** | 15 | 13 |
| PFM_monalysin-like | cd17904 | 31 | 229 | 300 | **230** | 66 | 4 | **212** | 85 | 3 | **225** | 62 | 13 | **225** | 60 | 15 | **215** | 76 | 9 |
| Mb-like | cd01040 | 384 | 239 | 300 | **287** | 11 | 2 | **284** | 14 | 2 | **287** | 8 | 5 | **284** | 11 | 5 | **283** | 14 | 3 |
| FYVE_like_SF | cd00065 | 392 | 266 | 300 | **266** | 25 | 9 | **262** | 23 | 15 | **264** | 19 | 17 | **269** | 18 | 13 | **263** | 24 | 13 |
| PFM_aerolysin_family | cd10140 | 65 | 270 | 300 | **283** | 17 | 0 | **276** | 24 | 0 | **261** | 36 | 3 | **260** | 38 | 2 | **252** | 46 | 2 |
| 7tmA_photoreceptors_insect | cd15079 | 35 | 301 | 300 | **236** | 44 | 20 | **248** | 29 | 23 | **243** | 34 | 23 | **244** | 33 | 23 | **240** | 37 | 23 |
| 7tmA_Melanopsin-like | cd15083 | 12 | 314 | 55 | **52** | 2 | 1 | **52** | 2 | 1 | **52** | 2 | 1 | **52** | 2 | 1 | **52** | 2 | 1 |
| ClyA_AhlB-like | cd22652 | 39 | 354 | 300 | **236** | 31 | 33 | **252** | 14 | 34 | **253** | 12 | 35 | **252** | 13 | 35 | **196** | 18 | 86 |
| 7tmA_Opsins_type2_animals | cd14969 | 71 | 400 | 300 | **296** | 2 | 2 | **290** | 8 | 2 | **297** | 1 | 2 | **297** | 1 | 2 | **295** | 2 | 3 |
| 7tm_GPCRs | cd14964 | 18 | 420 | 136 | **136** | 0 | 0 | **134** | 2 | 0 | **134** | 2 | 0 | **134** | 2 | 0 | **134** | 2 | 0 |
| 7tmA_Anaphylatoxin_R-like | cd14974 | 18 | 429 | 136 | **133** | 2 | 1 | **131** | 4 | 1 | **135** | 0 | 1 | **135** | 0 | 1 | **134** | 1 | 1 |
| 7tmA_Opioid_R-like | cd14970 | 16 | 458 | 105 | **102** | 3 | 0 | **99** | 6 | 0 | **99** | 5 | 1 | **103** | 0 | 2 | **103** | 1 | 1 |
| ClyA-like | cd21116 | 118 | 519 | 300 | **280** | 14 | 6 | **278** | 16 | 6 | **286** | 7 | 7 | **289** | 5 | 6 | **281** | 13 | 6 |
| FGGY_RBK_like | cd07768 | 6 | 537 | 10 | **10** | 0 | 0 | **9** | 1 | 0 | **10** | 0 | 0 | **10** | 0 | 0 | **10** | 0 | 0 |
| FGGY | cd00366 | 31 | 588 | 300 | **287** | 5 | 8 | **244** | 48 | 8 | **292** | 0 | 8 | **292** | 0 | 8 | **288** | 4 | 8 |
| FGGY_YpCarbK_like | cd07782 | 35 | 690 | 300 | **262** | 35 | 3 | **274** | 22 | 4 | **276** | 19 | 5 | **278** | 16 | 6 | **274** | 18 | 8 |
| 7tm_classA_rhodopsin-like | cd00637 | 405 | 808 | 300 | **299** | 1 | 0 | **294** | 6 | 0 | **299** | 1 | 0 | **299** | 1 | 0 | **299** | 1 | 0 |
| NBD_sugar-kinase_HSP70_actin | cd00012 | 125 | 1154 | 300 | **290** | 8 | 2 | **283** | 15 | 2 | **298** | 0 | 2 | **298** | 0 | 2 | **298** | 0 | 2 |
| 7tmA_amine_R-like | cd14967 | 78 | 1227 | 300 | **292** | 8 | 0 | **288** | 12 | 0 | **297** | 2 | 1 | **296** | 3 | 1 | **296** | 4 | 0 |
| **Total** | | | | | **7,767** | 1,485 | 1,336 | **7,625** | 1,512 | 1,451 | **7,771** | 1,207 | 1,610 | **7,798** | 1,227 | 1,563 | **7,425** | 1,457 | 1,706 |
| **Total %** | | | | | **0.73** | 0.14 | 0.13 | **0.72** | 0.14 | 0.14 | **0.73** | 0.11 | 0.15 | **0.74** | 0.12 | 0.15 | **0.70** | 0.14 | 0.16 |

#### 2 Best *E*-score

Supplementary Table 3: End-gap-free alignments: average distance for all E-scores and all five distances for six selected MSAs. Best results are shown in boldface.

| MSA | | | | | d_cc | | | | | |
|---|---|---|---|---|---|---|---|---|---|---|
| Conserved domain | Source | Proteins | Length | Samples | ProtT5 | ProtBert | ESM1b | ESM2 | ProtAlbert | XLNet |
| *Hb-alpha-like* | cd08927 | 39 | 142 | 300 | 0.000678 | **0.000510** | 0.000628 | 0.000555 | 0.000635 | - |
| *Globin-like* | cd01067 | 17 | 161 | 120 | **0.032387** | 0.049760 | 0.061074 | 0.035322 | 0.049610 | 0.06821 |
| *7tmA_photoreceptors_insect* | cd15079 | 35 | 301 | 300 | 0.001399 | 0.001319 | 0.001241 | **0.000967** | 0.001390 | 0.00211 |
| *7tm_GPCRs* | cd14964 | 18 | 420 | 136 | **0.022676** | 0.038388 | 0.031416 | 0.027154 | 0.033099 | 0.03873 |
| *FGGY_YpCarbK_like* | cd07782 | 35 | 690 | 300 | **0.001603** | 0.001793 | 0.002231 | 0.001730 | 0.002565 | - |
| *NBD_sugar-kinase_HSP70_actin* | cd00012 | 125 | 1154 | 300 | **0.033371** | 0.050246 | 0.046352 | 0.041707 | 0.049278 | - |

| MSA | | | | | d_d | | | | | | d_pos | | | | | |
|---|---|---|---|---|---|---|---|---|---|---|---|---|---|---|---|---|
| Conserved domain | Source | Proteins | Length | Samples | ProtT5 | ProtBert | ESM1b | ESM2 | ProtAlbert | XLNet | ProtT5 | ProtBert | ESM1b | ESM2 | ProtAlbert | XLNet |
| *Hb-alpha-like* | cd08927 | 39 | 142 | 300 | 0.000044 | 0.000037 | 0.000045 | **0.000037** | 0.000048 | - | 0.015327 | 0.012939 | 0.013942 | **0.012800** | 0.015381 | - |
| *Globin-like* | cd01067 | 17 | 161 | 120 | **0.006550** | 0.015153 | 0.017836 | 0.008764 | 0.011445 | 0.02027 | **0.370198** | 0.551089 | 0.613001 | 0.401669 | 0.544223 | 0.68865 |
| *7tmA_photoreceptors_insect* | cd15079 | 35 | 301 | 300 | 0.000228 | 0.000186 | 0.000157 | **0.000145** | 0.000184 | 0.00038 | 0.049456 | 0.046965 | 0.045116 | **0.035749** | 0.045387 | 0.07241 |
| *7tm_GPCRs* | cd14964 | 18 | 420 | 136 | **0.005327** | 0.018973 | 0.012929 | 0.011427 | 0.009080 | 0.01586 | **0.533216** | 0.686566 | 0.637672 | 0.556423 | 0.668073 | 0.69320 |
| *FGGY_YpCarbK_like* | cd07782 | 35 | 690 | 300 | **0.000224** | 0.000267 | 0.000438 | 0.000240 | 0.000385 | - | 0.063175 | 0.068498 | 0.078059 | **0.061100** | 0.071485 | - |
| *NBD_sugar-kinase_HSP70_actin* | cd00012 | 125 | 1154 | 300 | **0.018078** | 0.037083 | 0.035718 | 0.030538 | 0.034399 | - | **0.664004** | 0.795209 | 0.750176 | 0.709844 | 0.785006 | - |

| MSA | | | | | d_seq | | | | | | d_ssp | | | | | |
|---|---|---|---|---|---|---|---|---|---|---|---|---|---|---|---|---|
| Conserved domain | Source | Proteins | Length | Samples | ProtT5 | ProtBert | ESM1b | ESM2 | ProtAlbert | XLNet | ProtT5 | ProtBert | ESM1b | ESM2 | ProtAlbert | XLNet |
| *Hb-alpha-like* | cd08927 | 39 | 142 | 300 | 0.014910 | 0.012677 | 0.013799 | **0.012467** | 0.015167 | - | 0.015378 | 0.010555 | 0.014715 | **0.011833** | 0.016289 | - |
| *Globin-like* | cd01067 | 17 | 161 | 120 | **0.349277** | 0.544652 | 0.604901 | 0.388417 | 0.530987 | 0.68365 | **0.410540** | 0.617252 | 0.674917 | 0.452781 | 0.605535 | 0.75062 |
| *7tmA_photoreceptors_insect* | cd15079 | 35 | 301 | 300 | 0.047164 | 0.044023 | 0.042024 | **0.032658** | 0.043924 | 0.06998 | 0.065482 | 0.063650 | 0.063435 | **0.046808** | 0.062535 | 0.10647 |
| *7tm_GPCRs* | cd14964 | 18 | 420 | 136 | **0.480151** | 0.664946 | 0.601501 | 0.523269 | 0.628612 | 0.66842 | **0.609748** | 0.753075 | 0.712301 | 0.629215 | 0.738981 | 0.76306 |
| *FGGY_YpCarbK_like* | cd07782 | 35 | 690 | 300 | 0.050921 | 0.058269 | 0.067059 | **0.050394** | 0.060601 | - | 0.062732 | 0.068537 | 0.093033 | **0.060126** | 0.072615 | - |
| *NBD_sugar-kinase_HSP70_actin* | cd00012 | 125 | 1154 | 300 | **0.565443** | 0.748106 | 0.698232 | 0.656375 | 0.732283 | - | **0.688703** | 0.823625 | 0.781364 | 0.733834 | 0.814248 | - |

Supplementary Table 4: End-gap-free alignments: ProtT5-score vs ESM2-score for six selected MSAs. Best results are shown in boldface. Wilcoxon test P-values higher than .01 are shown in red.

| MSA | | | | | d_cc | | | d_d | | | d_pos | | | d_seq | | | d_ssp | | |
|---|---|---|---|---|---|---|---|---|---|---|---|---|---|---|---|---|---|---|---|
| Conserved domain | Source | Proteins | Length | Samples | ESM2 | ProtT5 | P-value | ESM2 | ProtT5 | P-value | ESM2 | ProtT5 | P-value | ESM2 | ProtT5 | P-value | ESM2 | ProtT5 | P-value |
| *Hb-alpha-like* | cd08927 | 39 | 142 | 300 | **0.000555** | 0.000678 | 9.32E-07 | **0.000037** | 0.000044 | 3.39E-05 | **0.012800** | 0.015327 | 1.65E-07 | **0.012467** | 0.014910 | 4.98E-07 | **0.011833** | 0.015378 | 4.29E-06 |
| *Globin-like* | cd01067 | 17 | 161 | 120 | 0.035322 | **0.032387** | 1.51E-02 | 0.008764 | **0.006550** | 2.26E-11 | 0.401669 | **0.370198** | 5.40E-03 | 0.388417 | **0.349277** | 1.26E-03 | 0.452781 | **0.410540** | 1.55E-03 |
| *7tmA_photoreceptors_insect* | cd15079 | 35 | 301 | 300 | **0.000967** | 0.001399 | 1.73E-29 | **0.000145** | 0.000228 | 9.58E-38 | **0.035749** | 0.049456 | 1.74E-39 | **0.032658** | 0.047164 | 8.10E-40 | **0.046808** | 0.065482 | 2.07E-38 |
| *7tm_GPCRs* | cd14964 | 18 | 420 | 136 | 0.027154 | **0.022676** | 6.58E-04 | 0.011427 | **0.005327** | 3.87E-10 | 0.556423 | **0.533216** | 3.83E-02 | 0.523269 | **0.480151** | 2.49E-04 | 0.629215 | **0.609748** | 1.04E-01 |
| *FGGY_YpCarbK_like* | cd07782 | 35 | 690 | 300 | 0.001730 | **0.001603** | 3.31E-03 | 0.000240 | **0.000224** | 1.66E-03 | **0.061100** | 0.063175 | 1.07E-03 | **0.050394** | 0.050921 | 4.10E-02 | **0.060126** | 0.062732 | 1.66E-08 |
| *NBD_sugar-kinase_HSP70_actin* | cd00012 | 125 | 1154 | 300 | 0.041707 | **0.033371** | 1.90E-18 | 0.030538 | **0.018078** | 1.85E-32 | 0.709844 | **0.664004** | 1.12E-16 | 0.656375 | **0.565443** | 1.34E-30 | 0.733834 | **0.688703** | 8.20E-15 |

### 3 Best BLOSUM matrix

Supplementary Table 5: End-gap-free alignments: average distance for all BLOSUM matrices and all five distances for six selected MSAs. Best results are shown in boldface.

| MSA | | | | | d_cc | | | | |
|---|---|---|---|---|---|---|---|---|---|
| Conserved domain | Source | Proteins | Length | Samples | BLOSUM45 | BLOSUM50 | BLOSUM62 | BLOSUM80 | BLOSUM90 |
| *Hb-alpha-like* | cd08927 | 39 | 142 | 300 | 0.000975 | 0.001074 | **0.000870** | 0.001106 | 0.001191 |
| *Globin-like* | cd01067 | 17 | 161 | 120 | **0.160539** | 0.163812 | 0.311554 | 0.384547 | 0.382977 |
| *7tmA_photoreceptors_insect* | cd15079 | 35 | 301 | 300 | 0.002451 | 0.002687 | **0.002305** | 0.002681 | 0.002876 |
| *7tm_GPCRs* | cd14964 | 18 | 420 | 136 | **0.098840** | 0.114038 | 0.291492 | 0.442796 | 0.444632 |
| *FGGY_YpCarbK_like* | cd07782 | 35 | 690 | 300 | 0.004856 | 0.004930 | **0.004560** | 0.004953 | 0.005228 |
| *NBD_sugar-kinase_HSP70_actin* | cd00012 | 125 | 1154 | 300 | **0.120524** | 0.124483 | 0.351562 | 0.436627 | 0.443355 |

| MSA | | | | | d_d | | | | | d_pos | | | | |
|---|---|---|---|---|---|---|---|---|---|---|---|---|---|---|
| Conserved domain | Source | Proteins | Length | Samples | BLOSUM45 | BLOSUM50 | BLOSUM62 | BLOSUM80 | BLOSUM90 | BLOSUM45 | BLOSUM50 | BLOSUM62 | BLOSUM80 | BLOSUM90 |
| *Hb-alpha-like* | cd08927 | 39 | 142 | 300 | 0.000228 | 0.000332 | **0.000172** | 0.000313 | 0.000557 | 0.022717 | 0.025696 | **0.021142** | 0.027009 | 0.029803 |
| *Globin-like* | cd01067 | 17 | 161 | 120 | **0.118599** | 0.124467 | 0.292858 | 0.376061 | 0.373764 | **0.806900** | 0.816397 | 0.846399 | 0.876281 | 0.881988 |
| *7tmA_photoreceptors_insect* | cd15079 | 35 | 301 | 300 | 0.001084 | 0.001500 | **0.000857** | 0.001484 | 0.002018 | 0.084005 | 0.090461 | **0.079186** | 0.089835 | 0.097210 |
| *7tm_GPCRs* | cd14964 | 18 | 420 | 136 | **0.071275** | 0.092714 | 0.291051 | 0.441464 | 0.444240 | **0.891019** | 0.902100 | 0.931473 | 0.972709 | 0.976773 |
| *FGGY_YpCarbK_like* | cd07782 | 35 | 690 | 300 | 0.001399 | 0.001774 | **0.001235** | 0.001788 | 0.002313 | 0.107904 | 0.111766 | **0.101768** | 0.111036 | 0.117510 |
| *NBD_sugar-kinase_HSP70_actin* | cd00012 | 125 | 1154 | 300 | **0.104434** | 0.109325 | 0.351568 | 0.435391 | 0.443592 | **0.903067** | 0.904574 | 0.939824 | 0.954303 | 0.956641 |

| MSA | | | | | d_seq | | | | | d_ssp | | | | |
|---|---|---|---|---|---|---|---|---|---|---|---|---|---|---|
| Conserved domain | Source | Proteins | Length | Samples | BLOSUM45 | BLOSUM50 | BLOSUM62 | BLOSUM80 | BLOSUM90 | BLOSUM45 | BLOSUM50 | BLOSUM62 | BLOSUM80 | BLOSUM90 |
| *Hb-alpha-like* | cd08927 | 39 | 142 | 300 | 0.022455 | 0.025446 | **0.020892** | 0.026795 | 0.029518 | 0.027156 | 0.031297 | **0.024384** | 0.031611 | 0.033224 |
| *Globin-like* | cd01067 | 17 | 161 | 120 | **0.775463** | 0.781519 | 0.786103 | 0.801414 | 0.806853 | **0.823240** | 0.830480 | 0.853761 | 0.879902 | 0.884695 |
| *7tmA_photoreceptors_insect* | cd15079 | 35 | 301 | 300 | 0.082066 | 0.088500 | **0.077444** | 0.087955 | 0.095279 | 0.114693 | 0.121905 | **0.107431** | 0.119600 | 0.127150 |
| *7tm_GPCRs* | cd14964 | 18 | 420 | 136 | 0.849745 | 0.856005 | **0.848253** | 0.867307 | 0.871364 | **0.912477** | 0.920913 | 0.942708 | 0.975132 | 0.978878 |
| *FGGY_YpCarbK_like* | cd07782 | 35 | 690 | 300 | 0.097829 | 0.101641 | **0.091791** | 0.100644 | 0.106800 | 0.121254 | 0.124388 | **0.112235** | 0.122356 | 0.128248 |
| *NBD_sugar-kinase_HSP70_actin* | cd00012 | 125 | 1154 | 300 | 0.825004 | 0.819877 | 0.761822 | **0.747700** | 0.748795 | 0.913651 | **0.913441** | 0.945692 | 0.957563 | 0.959222 |

Supplementary Table 6: End-gap-free alignments: BLOSUM45 vs BLOSUM62 matrices for six selected MSAs. Best results are shown in boldface. Wilcoxon test P-values higher than .01 are shown in red.

| MSA | | | | | d_cc | | | d_d | | | d_pos | | | d_seq | | | d_ssp | | |
|---|---|---|---|---|---|---|---|---|---|---|---|---|---|---|---|---|---|---|---|
| Conserved domain | Source | Proteins | Length | Samples | BLOSUM45 | BLOSUM62 | P-value | BLOSUM45 | BLOSUM62 | P-value | BLOSUM45 | BLOSUM62 | P-value | BLOSUM45 | BLOSUM62 | P-value | BLOSUM45 | BLOSUM62 | P-value |
| *Hb-alpha-like* | cd08927 | 39 | 142 | 300 | 0.000975 | **0.000870** | 5.35E-01 | 0.000228 | **0.000172** | 3.72E-01 | 0.022717 | **0.021142** | 6.94E-01 | 0.022455 | **0.020892** | 6.87E-01 | 0.027156 | **0.024384** | 3.24E-01 |
| *Globin-like* | cd01067 | 17 | 161 | 120 | **0.160539** | 0.311554 | 9.87E-16 | **0.118599** | 0.292858 | 2.21E-15 | **0.806900** | 0.846399 | 1.32E-06 | **0.775463** | 0.786103 | 9.58E-01 | **0.823240** | 0.853761 | 3.55E-06 |
| *7tmA_photoreceptors_insect* | cd15079 | 35 | 301 | 300 | 0.002451 | **0.002305** | 2.30E-05 | 0.001084 | **0.000857** | 1.64E-05 | 0.084005 | **0.079186** | 5.50E-06 | 0.082066 | **0.077444** | 2.12E-05 | 0.114693 | **0.107431** | 1.82E-06 |
| *7tm_GPCRs* | cd14964 | 18 | 420 | 136 | **0.098840** | 0.291492 | 8.75E-20 | **0.071275** | 0.291051 | 3.42E-19 | **0.891019** | 0.931473 | 1.82E-08 | 0.849745 | **0.848253** | 2.84E-02 | **0.912477** | 0.942708 | 5.50E-08 |
| *FGGY_YpCarbK_like* | cd07782 | 35 | 690 | 300 | 0.004856 | **0.004560** | 1.25E-09 | 0.001399 | **0.001235** | 2.73E-06 | 0.107904 | **0.101768** | 2.21E-12 | 0.097829 | **0.091791** | 1.07E-11 | 0.121254 | **0.112235** | 4.35E-12 |
| *NBD_sugar-kinase_HSP70_actin* | cd00012 | 125 | 1154 | 300 | **0.120524** | 0.351562 | 1.71E-46 | **0.104434** | 0.351568 | 2.38E-46 | **0.903067** | 0.939824 | 3.71E-24 | 0.825004 | **0.761822** | 4.53E-26 | **0.913651** | 0.945692 | 5.84E-24 |

### 4 Global alignment tests

Supplementary Table 7: Global alignments: ProtT5-score vs BLOSUM45 matrix, average distances for all five distances and all testing MSAs. Best results are shown in boldface. Wilcoxon test P-values higher than .01 are shown in red.

| MSA | | | | | d_cc | | | d_d | | | d_pos | | | d_seq | | | d_ssp | | |
|---|---|---|---|---|---|---|---|---|---|---|---|---|---|---|---|---|---|---|---|
| Conserved domain | Source | Proteins | Length | Samples | ProtT5 | BLOSUM45 | P-value | ProtT5 | BLOSUM45 | P-value | ProtT5 | BLOSUM45 | P-value | ProtT5 | BLOSUM45 | P-value | ProtT5 | BLOSUM45 | P-value |
| Bbox2_MID2_C-I | cd19823 | 8 | 40 | 21 | 0.000000 | 0.000000 | - | 0.000000 | 0.000000 | - | 0.000000 | 0.000000 | - | 0.000000 | 0.000000 | - | 0.000000 | 0.000000 | - |
| Bbox2_TRIM42_C-III | cd19782 | 9 | 40 | 28 | 0.003005 | **0.001694** | 6.79E-02 | 0.000174 | **0.000081** | 6.79E-02 | 0.016727 | **0.008590** | 6.79E-02 | 0.016727 | **0.008590** | 6.79E-02 | 0.020341 | **0.008104** | 6.79E-02 |
| Bbox2_MID | cd19758 | 10 | 40 | 36 | 0.001193 | **0.001006** | 4.31E-02 | 0.000361 | **0.000289** | 3.94E-02 | 0.033052 | **0.026723** | 3.94E-02 | 0.033052 | **0.026723** | 3.94E-02 | 0.044817 | **0.035434** | 3.94E-02 |
| Bbox2_MID1_C-I | cd19822 | 9 | 47 | 28 | 0.000000 | 0.000000 | - | 0.000000 | 0.000000 | - | 0.000000 | 0.000000 | - | 0.000000 | 0.000000 | - | 0.000000 | 0.000000 | - |
| Bbox_SF | cd00021 | 6 | 48 | 10 | **0.023063** | 0.031005 | 4.84E-01 | **0.011057** | 0.011504 | 8.89E-01 | **0.236113** | 0.246558 | 8.89E-01 | 0.234950 | **0.231937** | 8.66E-01 | 0.241883 | **0.240388** | 8.66E-01 |
| DEFL_defensin-like | cd21806 | 78 | 51 | 300 | 0.010128 | **0.008159** | 6.34E-07 | 0.002951 | **0.002898** | 2.65E-02 | 0.119022 | **0.111145** | 1.72E-02 | 0.104142 | **0.096748** | 1.62E-02 | 0.097772 | **0.097251** | 1.80E-01 |
| Bbox2_TRIM37_C-VIII | cd19779 | 25 | 52 | 276 | 0.004326 | **0.003154** | 6.05E-09 | 0.001326 | **0.000833** | 3.58E-14 | 0.050800 | **0.034616** | 1.25E-13 | 0.044965 | **0.026834** | 1.18E-14 | 0.037020 | **0.021853** | 3.59E-13 |
| Bbox2_TRIM9-like | cd19764 | 17 | 53 | 120 | 0.006689 | **0.004010** | 6.92E-05 | 0.001876 | **0.001368** | 1.08E-02 | 0.066398 | **0.054257** | 4.93E-03 | 0.052880 | **0.043590** | 3.37E-02 | 0.049004 | **0.043404** | 6.68E-02 |
| CBD_like | cd12204 | 41 | 61 | 300 | **0.010804** | 0.014363 | 6.95E-10 | **0.003781** | 0.005127 | 2.61E-08 | **0.152429** | 0.184270 | 1.11E-11 | **0.130087** | 0.166340 | 1.45E-13 | **0.151414** | 0.185038 | 2.02E-06 |
| Bbox2 | cd19756 | 127 | 65 | 300 | **0.011511** | 0.012688 | 9.71E-01 | **0.002942** | 0.003390 | 2.56E-01 | 0.134007 | **0.130947** | 3.43E-03 | 0.127192 | **0.124068** | 9.75E-03 | 0.143595 | **0.140827** | 8.86E-03 |
| ChtBD1 | cd00035 | 32 | 67 | 300 | **0.012408** | 0.014017 | 1.23E-02 | **0.003179** | 0.004553 | 2.66E-05 | **0.157288** | 0.176369 | 7.37E-04 | **0.128726** | 0.147971 | 2.05E-04 | **0.124246** | 0.149246 | 2.25E-04 |
| Bbox1_CYLD | cd19816 | 27 | 68 | 325 | **0.013647** | 0.017508 | 1.81E-05 | **0.004508** | 0.005065 | 9.85E-01 | **0.154754** | 0.169441 | 2.77E-03 | **0.146010** | 0.160007 | 2.65E-03 | **0.164416** | 0.186957 | 4.62E-04 |
| KAZAL_FS | cd00104 | 273 | 74 | 300 | **0.013922** | 0.028184 | 3.84E-22 | **0.003314** | 0.008707 | 1.79E-22 | **0.145909** | 0.227736 | 2.31E-20 | **0.136510** | 0.218210 | 5.27E-20 | **0.158976** | 0.250152 | 5.22E-17 |
| bHLH_SF | cd00083 | 79 | 75 | 300 | **0.007183** | 0.014852 | 1.27E-18 | **0.002838** | 0.006743 | 1.51E-17 | **0.102140** | 0.163804 | 4.04E-16 | **0.079882** | 0.149265 | 3.48E-18 | **0.079658** | 0.171462 | 1.25E-18 |
| CD_CSD | cd00024 | 522 | 98 | 300 | **0.009396** | 0.013120 | 4.39E-13 | **0.001655** | 0.003690 | 7.34E-25 | **0.110263** | 0.157445 | 2.75E-19 | **0.100134** | 0.149348 | 1.52E-20 | **0.105011** | 0.177089 | 4.61E-22 |
| C1 | cd00029 | 281 | 99 | 300 | **0.013681** | 0.018442 | 1.07E-10 | **0.003766** | 0.005289 | 1.61E-06 | **0.165077** | 0.204748 | 3.38E-09 | **0.156099** | 0.193662 | 5.46E-08 | **0.174556** | 0.226230 | 2.90E-08 |
| TrHb | cd14756 | 8 | 130 | 21 | **0.004520** | 0.025456 | 1.32E-04 | **0.000995** | 0.005879 | 2.14E-04 | **0.092407** | 0.290605 | 1.32E-04 | **0.081824** | 0.283148 | 1.32E-04 | **0.089073** | 0.357702 | 1.32E-04 |
| Hb | cd14765 | 15 | 138 | 91 | **0.001490** | 0.002370 | 8.92E-06 | **0.000204** | 0.000583 | 9.73E-09 | **0.044115** | 0.069436 | 4.91E-08 | **0.034136** | 0.062930 | 5.16E-10 | **0.041652** | 0.076554 | 5.03E-09 |
| SH2_STAT5 | cd10376 | 5 | 140 | 6 | **0.001355** | 0.002005 | 4.58E-01 | **0.000121** | 0.000146 | 4.58E-01 | 0.020036 | **0.017497** | 4.58E-01 | 0.018824 | **0.016285** | 4.58E-01 | 0.015888 | **0.011003** | 4.58E-01 |
| Hb-beta-like | cd08925 | 27 | 140 | 300 | **0.001107** | 0.001862 | 4.23E-06 | **0.000179** | 0.000499 | 2.00E-09 | **0.023368** | 0.034923 | 7.42E-09 | **0.021182** | 0.032321 | 5.40E-08 | **0.020466** | 0.038795 | 4.80E-10 |
| Hb-alpha-like | cd08927 | 39 | 142 | 300 | **0.000678** | 0.000937 | 5.06E-03 | **0.000044** | 0.000200 | 2.86E-04 | **0.015327** | 0.021191 | 6.51E-03 | **0.014910** | 0.020930 | 5.40E-03 | **0.015378** | 0.025827 | 9.99E-04 |
| SH2_STAT5a | cd10421 | 10 | 145 | 36 | **0.001055** | 0.001406 | 1.16E-01 | **0.000129** | 0.000151 | 1.01E-01 | **0.014668** | 0.016317 | 7.81E-02 | **0.011324** | 0.013092 | 7.81E-02 | **0.007680** | 0.010716 | 7.81E-02 |
| Mb | cd08926 | 9 | 149 | 28 | 0.000681 | **0.000653** | 3.79E-01 | **0.000049** | 0.000072 | 2.53E-02 | **0.014150** | 0.017551 | 1.95E-01 | **0.013665** | 0.017551 | 8.69E-02 | **0.010430** | 0.018042 | 2.59E-02 |
| GS_GGDEF_2 | cd14759 | 26 | 152 | 300 | **0.001836** | 0.007858 | 2.33E-28 | **0.000270** | 0.002425 | 1.16E-32 | **0.027214** | 0.122792 | 2.15E-35 | **0.026258** | 0.121768 | 4.57E-35 | **0.034206** | 0.170564 | 1.52E-34 |
| Globin-like | cd01067 | 17 | 161 | 120 | **0.031733** | 0.064032 | 1.21E-17 | **0.006543** | 0.026334 | 1.26E-20 | **0.368170** | 0.659491 | 1.23E-20 | **0.347358** | 0.642125 | 7.01E-21 | **0.408049** | 0.710038 | 7.19E-21 |
| FHb-globin | cd08922 | 47 | 162 | 300 | **0.001234** | 0.002119 | 3.83E-12 | **0.000146** | 0.000537 | 1.76E-25 | **0.025158** | 0.038901 | 6.24E-21 | **0.019062** | 0.034087 | 3.90E-22 | **0.018162** | 0.035502 | 3.70E-20 |
| PFM_HFR-2-like | cd20216 | 90 | 174 | 300 | 0.006101 | **0.005901** | 7.67E-01 | **0.001517** | 0.001679 | 1.41E-04 | **0.098214** | 0.113686 | 2.92E-06 | **0.097322** | 0.112365 | 3.91E-06 | **0.137490** | 0.158859 | 5.70E-06 |
| SH2_STAT_family | cd09919 | 67 | 206 | 300 | **0.019460** | 0.021276 | 1.34E-03 | **0.005518** | 0.006777 | 1.66E-03 | **0.203658** | 0.252028 | 2.34E-19 | **0.178629** | 0.224965 | 1.22E-16 | **0.208088** | 0.264876 | 7.82E-18 |
| PBP-like | cd08919 | 31 | 213 | 300 | **0.011902** | 0.023623 | 3.30E-37 | **0.003315** | 0.007036 | 3.67E-25 | **0.189161** | 0.291599 | 3.08E-36 | **0.176688** | 0.280838 | 9.80E-36 | **0.215297** | 0.342865 | 1.09E-35 |
| SH2 | cd00173 | 352 | 214 | 300 | **0.033438** | 0.045111 | 7.33E-22 | **0.009190** | 0.013590 | 2.27E-20 | **0.299843** | 0.410207 | 1.17E-30 | **0.275217** | 0.388500 | 4.20E-30 | **0.327982** | 0.463043 | 2.24E-29 |
| Globin_sensor | cd01068 | 193 | 223 | 300 | **0.004459** | 0.016223 | 1.49E-44 | **0.000583** | 0.004483 | 6.65E-47 | **0.068155** | 0.208243 | 3.84E-46 | **0.051818** | 0.197550 | 1.50E-46 | **0.061683** | 0.250642 | 5.03E-46 |
| PFM_monalysin-like | cd17904 | 31 | 229 | 300 | **0.013546** | 0.015735 | 6.40E-07 | **0.005046** | 0.005114 | 3.71E-06 | **0.263529** | 0.286888 | 1.36E-07 | **0.261302** | 0.284632 | 2.12E-07 | **0.346110** | 0.372784 | 6.82E-07 |
| Mb-like | cd01040 | 384 | 239 | 300 | **0.008814** | 0.025598 | 3.67E-45 | **0.002241** | 0.009006 | 5.63E-43 | **0.166566** | 0.359807 | 1.85E-46 | **0.141470** | 0.346156 | 7.23E-47 | **0.163770** | 0.416792 | 9.51E-47 |
| FYVE_like_SF | cd00065 | 392 | 266 | 300 | **0.012167** | 0.031618 | 4.97E-41 | **0.005300** | 0.013507 | 1.80E-36 | **0.181647** | 0.292302 | 6.76E-39 | **0.108226** | 0.224827 | 5.07E-41 | **0.123816** | 0.259494 | 5.38E-39 |
| PFM_aerolysin_family | cd10140 | 65 | 270 | 300 | **0.045061** | 0.056760 | 4.10E-26 | **0.010532** | 0.019177 | 3.29E-29 | **0.642903** | 0.721366 | 3.55E-21 | **0.608259** | 0.693639 | 3.82E-23 | **0.719626** | 0.787133 | 4.20E-18 |
| 7tmA_photoreceptors_insect | cd15079 | 35 | 301 | 300 | **0.001399** | 0.002368 | 1.55E-31 | **0.000228** | 0.001052 | 5.02E-34 | **0.049456** | 0.075499 | 7.19E-31 | **0.047164** | 0.073559 | 2.90E-31 | **0.065482** | 0.105396 | 1.89E-31 |
| 7tmA_Melanopsin-like | cd15083 | 12 | 314 | 55 | **0.000879** | 0.003824 | 8.74E-10 | **0.000131** | 0.001804 | 8.74E-10 | **0.043049** | 0.109751 | 1.36E-09 | **0.034671** | 0.102987 | 1.09E-09 | **0.038653** | 0.138049 | 1.52E-09 |
| ClyA_AhlB-like | cd22652 | 39 | 354 | 300 | **0.001996** | 0.002498 | 6.75E-08 | **0.000187** | 0.000827 | 1.90E-20 | **0.033045** | 0.063414 | 2.11E-23 | **0.031856** | 0.061980 | 1.40E-22 | **0.042178** | 0.088833 | 6.11E-23 |
| 7tmA_Opsins_type2_animals | cd14969 | 71 | 400 | 300 | **0.001943** | 0.007413 | 1.31E-49 | **0.000562** | 0.003557 | 7.58E-48 | **0.069111** | 0.207764 | 2.95E-50 | **0.056882** | 0.197296 | 2.92E-50 | **0.067174** | 0.260493 | 3.13E-50 |
| 7tm_GPCRs | cd14964 | 18 | 420 | 136 | **0.021977** | 0.046764 | 1.22E-22 | **0.005309** | 0.022659 | 1.04E-23 | **0.530298** | 0.801367 | 5.48E-23 | **0.477233** | 0.766608 | 2.29E-23 | **0.607210** | 0.845613 | 1.24E-22 |
| 7tmA_Anaphylatoxin_R-like | cd14974 | 18 | 429 | 136 | **0.003833** | 0.008599 | 2.66E-22 | **0.001186** | 0.003628 | 1.11E-21 | **0.109582** | 0.198233 | 2.81E-23 | **0.080050** | 0.172128 | 3.84E-23 | **0.101411** | 0.229464 | 3.08E-23 |
| 7tmA_Opioid_R-like | cd14970 | 16 | 458 | 105 | **0.002042** | 0.006304 | 7.78E-19 | **0.000973** | 0.002897 | 2.67E-17 | **0.117470** | 0.191432 | 3.60E-18 | **0.047128** | 0.126863 | 1.25E-18 | **0.048755** | 0.165182 | 8.78E-19 |
| ClyA-like | cd21116 | 118 | 519 | 300 | **0.022690** | 0.037783 | 2.74E-34 | **0.013635** | 0.024122 | 2.53E-30 | **0.429975** | 0.624160 | 1.20E-46 | **0.376312** | 0.589775 | 2.56E-47 | **0.462390** | 0.679178 | 4.11E-46 |
| FGGY_RBK_like | cd07768 | 6 | 537 | 10 | **0.005103** | 0.007279 | 1.95E-03 | **0.001284** | 0.003284 | 1.95E-02 | **0.227322** | 0.311245 | 1.95E-03 | **0.206354** | 0.299167 | 1.95E-03 | **0.282124** | 0.378646 | 1.95E-03 |
| FGGY | cd00366 | 31 | 588 | 300 | **0.004575** | 0.009960 | 1.05E-48 | **0.002272** | 0.004811 | 1.89E-34 | **0.188870** | 0.352219 | 2.74E-49 | **0.178342** | 0.344821 | 1.84E-49 | **0.238063** | 0.429620 | 2.43E-49 |
| FGGY_YpCarbK_like | cd07782 | 35 | 690 | 300 | **0.001603** | 0.002904 | 3.06E-37 | **0.000224** | 0.001088 | 7.15E-45 | **0.063175** | 0.101153 | 7.85E-48 | **0.050921** | 0.090108 | 1.68E-47 | **0.062732** | 0.115266 | 3.31E-47 |
| 7tm_classA_rhodopsin-like | cd00637 | 405 | 808 | 300 | **0.006051** | 0.019284 | 6.46E-51 | **0.002059** | 0.007700 | 3.58E-49 | **0.188327** | 0.417532 | 6.79E-51 | **0.154874** | 0.393975 | 6.79E-51 | **0.201403** | 0.496238 | 7.00E-51 |
| NBD_sugar-kinase_HSP70_actin | cd00012 | 125 | 1154 | 300 | **0.032877** | 0.052502 | 1.73E-44 | **0.018053** | 0.036214 | 3.38E-41 | **0.663322** | 0.838091 | 1.47E-50 | **0.564686** | 0.776516 | 1.29E-50 | **0.687956** | 0.860119 | 1.49E-50 |
| 7tmA_amine_R-like | cd14967 | 78 | 1227 | 300 | **0.004752** | 0.012643 | 6.34E-50 | **0.002395** | 0.008252 | 1.50E-46 | **0.184162** | 0.326660 | 1.22E-50 | **0.106436** | 0.252929 | 1.15E-50 | **0.137242** | 0.326527 | 8.22E-51 |

### 5 Matrices heatmaps

Supplementary Table 8: All five BLOSUM matrices and all twelve $E$-score matrices for the *NBD_sugar-kinase_HSP70_actin* MSA.

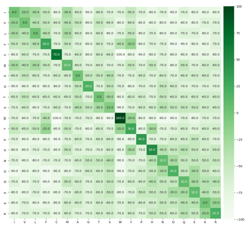

BLOSUM45

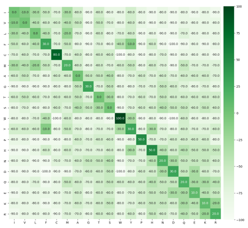

BLOSUM50

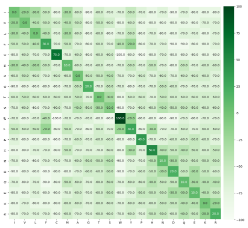

BLOSUM45

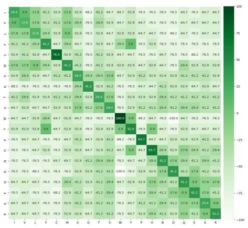

BLOSUM80

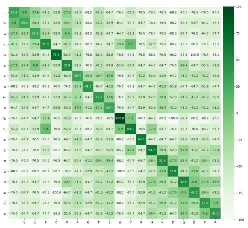

BLOSUM90

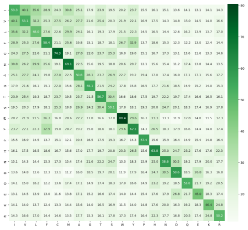

ProtT5 (A)

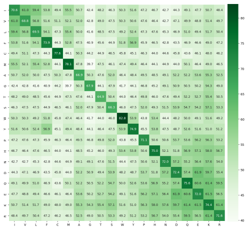

ESM2 (A)

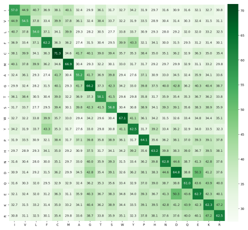

ProtBert (A)

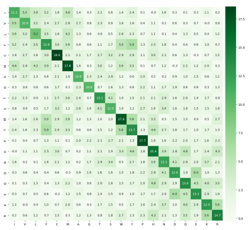

ProtT5 (U)

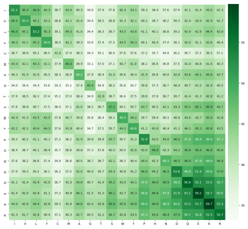

ESM2 (U)

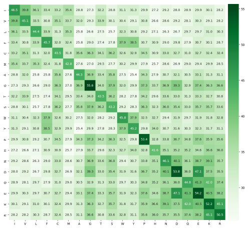

ProtBert (U)

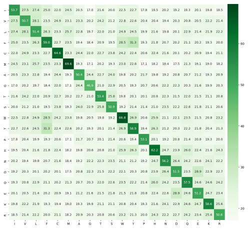

ProtAlbert (A)

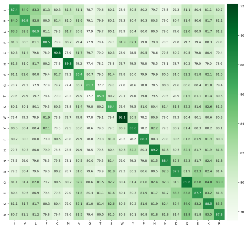

ESM-1b (A)

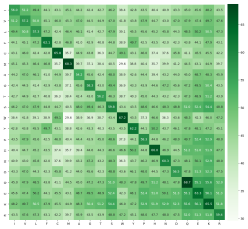

ProtXLNet (A)

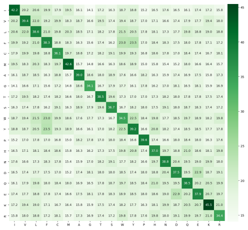

ProtAlbert (U)

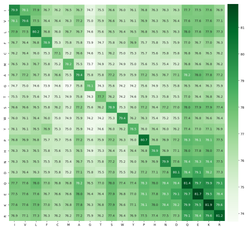

ESM-1b (U)

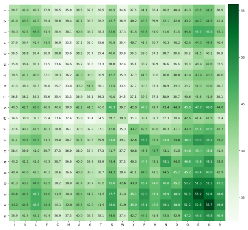

ProtXLNet (U)
